# A reusable neural approach to recombination mapping for model and non-model species

**DOI:** 10.64898/2026.08.20.746066

**Authors:** Kevin Korfmann, Neda Rahnamae, Sara Mathieson

## Abstract

Pedigree and crossing experiments can measure crossovers directly and provide the gold standard for recombination mapping, but their cost restricts fine-scale recombination mapping to only a few species. Patterns of linkage disequilibrium (LD) provide an alternative statistical approach for inferring variation in recombination along the genome. LD, however, is confounded by many evolutionary factors, such as demographic changes, life-history traits, and genomic structural variation. We present `fastrho`, a state-space neural-network estimator trained across a range of simulation-based priors. In simulated bottleneck and expansion scenarios, the fixed checkpoint recovered local map shape without target-specific retraining; comparisons with `pyrho` used lookup tables constructed under the simulation-generating history. We further evaluated generalizability across multiple species and, to account for additional confounders not represented in the initial training data, designed specialized models for inference in selfing plants, structured *Arabis* populations, and large-*N_e_* malaria-vector populations. A major biological application of the mosquito model was the construction of a five-arm recombination atlas spanning 13 Ag3 populations, providing a detailed view of recombination-rate variation across the dataset. Recombination maps inferred from Ag3 pedigrees provided independent, coarse-scale support for this atlas. Finally, analyses of resistance loci and redpoll bird supergenes demonstrate how selection and structural variation influence LD. Throughout our study, we use experimental maps for independent validation. Together, our results establish `fastrho` as a flexible framework for robust recombination mapping across diverse biological systems.

---

Recombination is responsible for shuffling and reconnecting haplotypes across generations according to the local crossover rate. This process determines how quickly linkage disequilibrium (LD), which refers to the local association of alleles, can decay along the genome. It is thus useful to detect fine-scale changes in recombination rates for studying adaptation, changes in genomic architecture, or the transmission of linked sites. For example, inversions, supergenes, or loci under selection create linkage patterns that can look like decreased recombination (1, 2). Much theory about the evolutionary importance of recombination has been developed over the years (3, 4).

LD is not affected by recombination alone; on the contrary, there are several confounders at work, including demographic effects, which directly affect LD and are consequently used to estimate population sizes (5). The relationship between genealogies and LD can also depend on sampling history (6). Other biological processes, such as selfing, constrain the effective number of recombination events, while structured populations and the resulting correlations among local genealogies distort the LD decay pattern (7). Furthermore, changes in the offspring distribution also have consequences for the correlation structure throughout the genome, resulting in a spiky LD decay pattern with many long-range structures (8). Returning to the effect of population size, in LD studies one usually infers the population-scaled value *ρ* = 4*N*_*e*_*r*, which means that any error in estimating *N*_*e*_ will propagate to the absolute rate. Thus, inferred hot or cold recombination spots can reflect not only the rate itself, but also the broader evolutionary process. Last but not least, this motivates the application of high-parameter simulation-based deep-learning models, which can account for the interaction of layered phenomena when these are represented during training.

Methods based on likelihood calculations, such as LDhat, LD-helmet, and `pyrho`, have enabled LD-based mapping in many systems, but their lookup tables and demographic corrections typically rely on prespecified models or assume simplifying assumption like constant population sizes (9–13). At biobank scale, FastRecomb uses PBWT-based haplotype matching to estimate population-specific maps with computation that scales linearly in the numbers of sites and individuals (14). Recent work in the coppery titi monkey illustrates the value of fine-scale re-combination maps for emerging model species (15). Estimators using simulations, such as FastEPRR, defiNETti, and `ReLERNN`, have expanded the methodological catalogue, but their reliability is contingent upon the ability of the simulations to approximate the target population; `ReLERNN` involves explicit simulation and model training for each target dataset (16–18). This requirement can become prohibitive when many populations or species are studied, because each application requires a separate simulation and training cycle based on a chosen demographic model. However, some robustness to demographic misspecification has been reported in the literature. Pedigree and cross maps provide the gold standard, but producing them in useful numbers and at sufficient resolution can entail dealing with hundreds of individuals. Sequencing-based approaches such as ReMIX instead use linked-read sequencing of gamete DNA from a single individual to detect crossover events (19). A complementary approach on the theoretical side is iSMC, which extends the sequentially Markovian coalescent to jointly infer demography and variation in recombination from as little as one unphased diploid genome, enabling applications to sample-limited and ancient genomes (20).

We address the problem of complex, layered evolutionary forces interacting with each other through amortization; in other words, the cost of learning the general simulation prior is paid during training, after which the same estimator may be used for other datasets. The base prior of `fastrho` uses outbred populations and is trained using several demographic categories (constant, sawtooth, island, and bottleneck), while checkpoints for special biological cases, including selfing, population structure, and large populations, are added individually. `fastrho` predicts the population-scaled rate *ρ* without requiring *N*_*e*_ as input; the absolute rate is calculated afterward as *r* = *ρ*/(4*N*_*e*_). For the simulation case, we employ the generating current or baseline *N*_*e*_, rather than any kind of average taken over the entire demographic history. On the other hand, in the real data case, the user can either specify an external *N*_*e*_ value or employ an auxiliary estimation that is based on the diversity features, and conditional upon the mutation rate, which makes it different from knowing the entire demographic history. The base model of `fastrho` is not trained on any species group; the species and maps from `stdpopsim` are instead used as realistic holdout transfer experiments.

The following real data experiments provide a stepwise evaluation of this idea. We use a ten-species screen and a multi-chromosome analysis to test whether `fastrho` can be reused across datasets. We further check the canid bottleneck, to analyze its robustness to demographic change, while the selfing and structured *Arabis* populations demonstrate why specialized checkpoints are needed to extend its range of application (21). The *N*_*e*_-specialized checkpoint is then used to construct a five-arm recombination atlas spanning 13 populations from the MalariaGEN Ag3 resource (22). The absolute rates in this atlas are conditional on diversity-informed *N*_*e*_ estimates and an assumed mutation rate of 3.5 × 10^−9^ per bp per generation. Recombination maps obtained from experimental crosses and pedigrees provide independent, coarse-scale support for the spatial variation in the atlas, but not for its absolute rate scale. Finally, we use the `fastrho` framework to study LD patterns associated with recombination, genomic architecture, and selection, focusing on the 2La inversion in Ag3, insecticide-resistance regions, and the redpoll supergene (23, 24). Thus, we aim to show the robust estimation of recombination from LD while using experimental recombination maps for independent comparison.

## Results

### A broad-prior LD estimator transfers across demographic scenarios

The key innovation of the implementation is that the general `fastrho` checkpoint can be learned once and then repeatedly applied to different demographic setups and real data without necessarily retraining it for each target. This model works on the level of segregating sites and makes use of physical distance, allele frequencies, multi-scale linkage disequilibrium, adjacent pair haplotypes, and local diversity. We summarize the model architecture, along inputs, and uncertainty-aware outputs in Fig. 1 and more detailed in the methods and SI Appendix.

**Figure 1.**
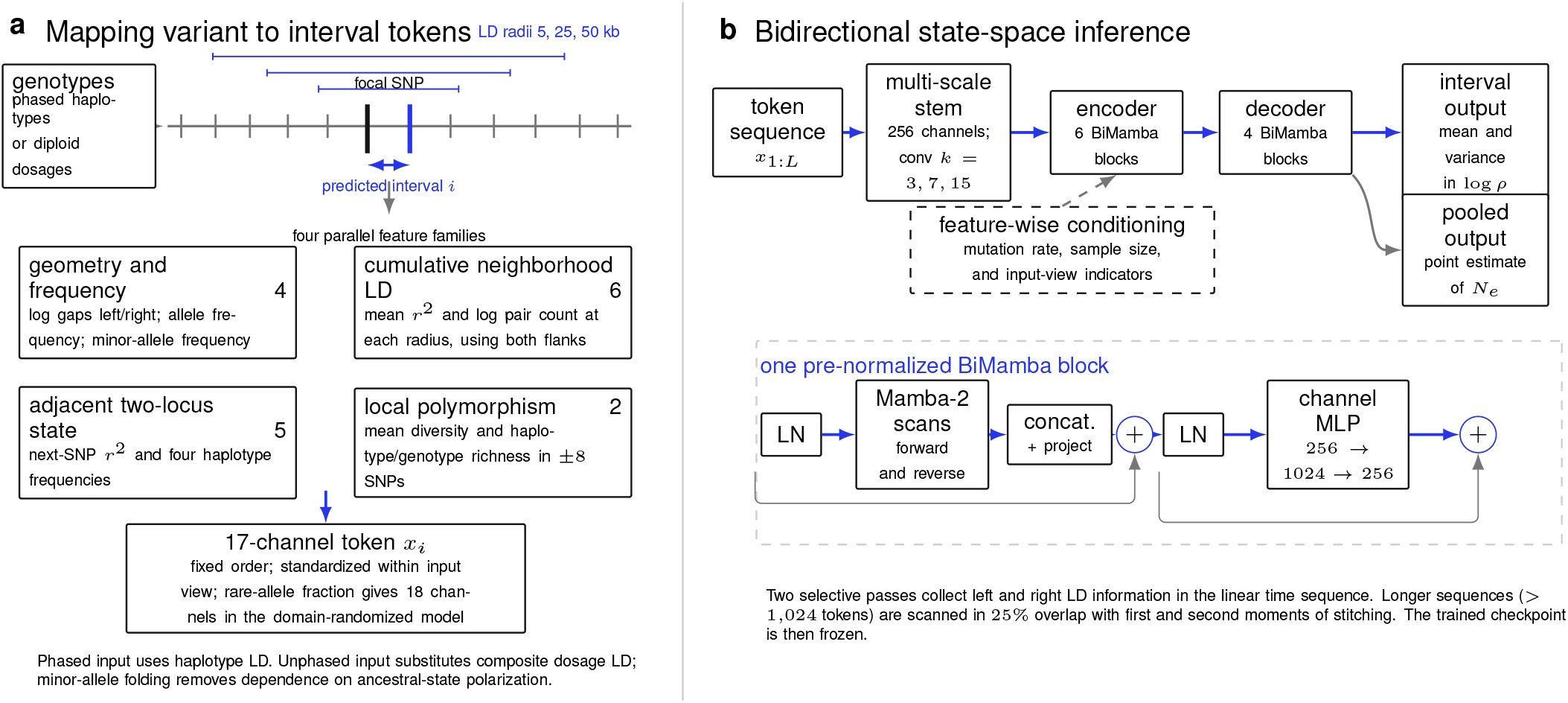
Input data and sequence modeling. (a) Every focal SNP is represented by a 17-value input vector describing the interval on its right. For unphased or unpolarized data, features that require phase or knowledge of the ancestral allele are replaced by features that do not. The model trained across these different input representations adds an eighteenth feature: the local proportion of SNPs with a minor-allele count of two or less. (b) The multi-scale input layer provides inputs to six encoder and four decoder BiMamba layers. The model predicts a Gaussian distribution of log *ρ* for each interval and also predicts a point estimate of *N*_*e*_.

We evaluated six demographic scenarios at a resolution of 25 kb (25). These consisted of three demographic models (constant, bottleneck, and expansion) and published recombination profiles based on deCODE, HapMap, and dog pedigree maps (26–28). `fastrho` was not fitted to a target demographic model, whereas `pyrho` used the generating history in the paired bottleneck and expansion comparisons. At 25 kb, Pearson correlations with the true maps ranged from 0.604 to 0.886 for `fastrho` and from 0.444 to 0.840 for `pyrho` (Fig. 2A, upper subpanel; SI Appendix, Tables S6 and S8). Because `ReLERNN` is a window-averaged estimator, we evaluated all three methods separately at the `ReLERNN` window scale. For each scenario, 20 prespecified 10-Mb test regions were simulated with a single recombination rate per region, equal to the length-weighted mean of the corresponding source map; the true rate and all three predictions were then averaged within each complete `ReLERNN` output window. Across the six scenarios, window-matched correlations ranged from 0.942 to 0.978 for `fastrho`, from 0.943 to 0.990 for `pyrho`, and from 0.847 to 0.946 for `ReLERNN` (Fig. 2A, lower subpanel; SI Appendix, Table S7).

**Figure 2.**
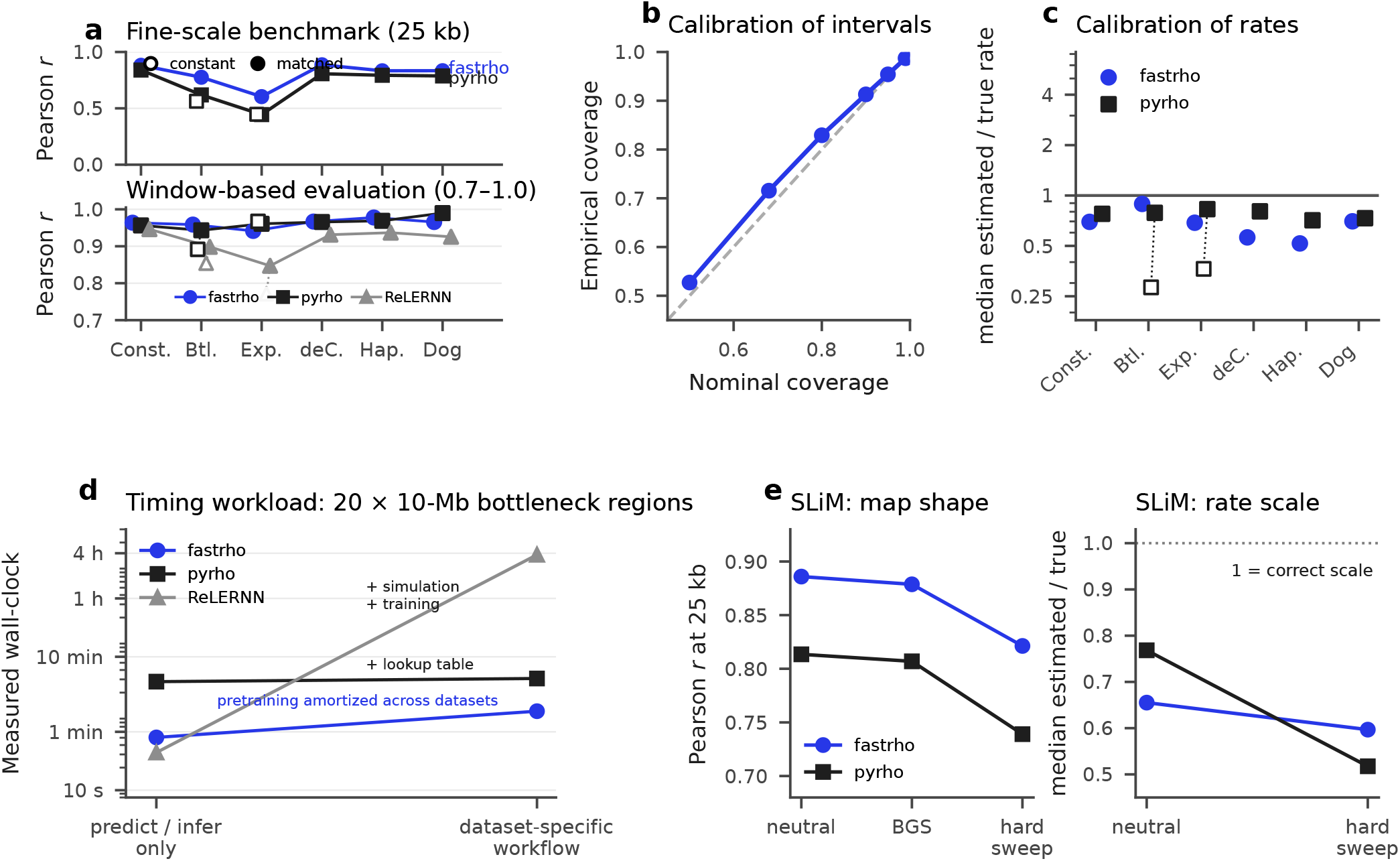
Simulation qualification. All benchmark regions used 20 sampled haplotypes; the fine-scale regions were 2 Mb. For human-like cases, parameters *N*_*e*_ = 10,000 and 1.5 × 10^−8^ mutations/bp/generation were used; for dogs, *N*_*e*_ = 13,000 and 4 × 10^−9^ mutations/bp/generation were used. (a) Pearson correlation for six scenarios. The upper subpanel compares `fastrho` and `pyrho` on heterogeneous maps at 25 kb. The lower subpanel, shown on an expanded 0.70–1.00 axis, compares all three methods on 20 prespecified 10-Mb constant-rate regions per scenario, scored within complete `ReLERNN` output windows. `ReLERNN` used phased input with at most 1,750 segregating sites per window (`--maxSites 1750`); median output-window widths were 0.74, 1.65, 2.85, 0.73, 0.74, and 2.28 Mb for constant, bottleneck, expansion, deCODE, HapMap, and dog, respectively. Filled symbols denote the generating demography; open symbols denote constant-history controls for bottleneck and expansion. `fastrho` uses the same fixed checkpoint throughout. (b) Observed versus expected interval coverage for 365,280 hidden intervals with known simulated *N*_*e*_ provided. (c) Median estimated-to-true rate at 25 kb; one indicates correct scale. (d) Measured wall-clock on the same 20 phased 10-Mb bottleneck regions used in the lower subpanel of (a). The left endpoint is prediction or inference only; the right is the dataset-specific workflow: checkpoint loading plus prediction for `fastrho`, lookup-table construction plus inference for `pyrho`, and simulation, 100-epoch training, and prediction for `ReLERNN`. `fastrho`’s upstream simulation and pretraining are amortized across datasets. (e) Independent 40-region SLiM simulations of map shape under neutrality, background selection, and a hard sweep, and of rate scale under neutrality and the sweep.

In the window-based evaluation, matching the generating history changed the window-matched correlation from 0.892 to 0.943 for `pyrho` and from 0.854 to 0.898 for `ReLERNN` under the bottleneck. Under expansion, the corresponding constant-history and matched-history values were 0.967 and 0.961 for `pyrho` and 0.772 and 0.847 for `ReLERNN`. These deliberately misspecified controls used the identical 20 test regions and `ReLERNN` output windows; they were fixed before scoring and are shown as open symbols in Fig. 2A.

Across 365,280 held-out SNP intervals, nominal 95% intervals covered the true recombination rate 95.4% of the time when the simulated *N*_*e*_ was supplied (Fig. 2B). This result measures mean interval-level calibration across the held-out simulation mixture conditional on the specified *N*_*e*_; it does not include uncertainty in the conversion from *ρ* to an absolute rate. Absolute-rate calibration varied among scenarios despite recovery of relative map shape (Fig. 2A,C).

Panel D uses the same 20 phased 10-Mb bottleneck regions as the window-based accuracy benchmark. Prediction or inference alone took 50.5 s for `fastrho`, 279.8 s for `pyrho`, and 31.8 s for `ReLERNN`. The dataset-specific workflows took 113.0 s, 308.3 s, and 13,729.6 s (3.81 h), respectively: `fastrho` included checkpoint loading and prediction, `pyrho` added lookup-table construction, and `ReLERNN` added simulation and 100-epoch training (Fig. 2D). `fastrho`’s upstream simulation and pretraining are one-time costs amortized across datasets. In a separate SLiM analysis, correlations under neutrality, background selection, and a hard sweep were 0.886, 0.879, and 0.821 for `fastrho` and 0.813, 0.807, and 0.739 for `pyrho`, respectively (Fig. 2E). The estimated-to-true rate ratio changed from 0.655 under neutrality to 0.597 under the hard sweep for `fastrho`, and from 0.768 to 0.517 for `pyrho`. Both methods retained map-shape information, but linked selection biased the inferred scale.

In a separate gene-conversion experiment, we applied the fixed recombination-only checkpoint to data simulated from known crossover maps. Without gene conversion, the 25-kb correlation with the map was *r* = 0.873. At a gene-conversion initiation rate four times the mean recombination rate, the correlation was *r* = 0.870 for 100-bp mean tracts and *r* = 0.678 for 1,000-bp mean tracts. Under the 1,000-bp condition, the inferred mean rate was 2.89-fold higher than its paired no-conversion value. The distortion therefore depended jointly on initiation rate and tract length (SI Appendix, Fig. S1).

### The base checkpoint transfers across species and chromosomes

We next applied the fixed broad checkpoint to ten public datasets spanning mammals, insects and plants; accessions and sources are reported in the SI Appendix. The display uses one representative chromosome per species, except that the *Arabidopsis* result is averaged across five chromosomes. Three species could be compared with an external map. For the remaining seven, split-sample correlations ranged from 0.59 to 0.94 and agreement with `pyrho` ranged from 0.50 to 0.82.

To test whether the representative tracks reflected isolated chromosomes, we expanded three panels further, to include four autosomal arms in *Drosophila melanogaster* and five chromosomes each in the jewel wasp *Nasonia vitripennis* and European aspen *Populus tremula*. At 100 kb, correlation with the independent Comeron map in *Drosophila* had a chromosome mean of 0.606 (chromosome-bootstrap 95% interval 0.559–0.651). The five-chromosome mean split-sample Spearman correlations were 0.713 (0.640–0.786) in jewel wasp and 0.898 (0.874–0.924) in aspen. Together with the existing five-chromosome *A. thaliana* comparison and the five-chromosome canid analysis below, this follow-up tests chromosome-to-chromosome replication in five species. The external-map values measure accuracy at the stated resolution, whereas the split-sample values measure repeatability only (SI Appendix, Fig. S4).

This portability benchmark proved a reusable across typical outbred datasets. We next asked how far specialization could extend that baseline under recent bottlenecks, selfing and population structure, three settings in which LD departs from the broad model’s default operating conditions.

### Severe bottlenecks limit recoverable LD signal

Severe bottlenecks may reduce the amount of information about recombination retained in LD, but a closely related population that has experienced less severe bottlenecks may retain more signal. We explored this possibility by testing whether wolf data would enhance recovery of the Campbell domestic-dog pedigree map (28, 29). For the first 40 Mb of chromosome 1, correlations measured within 100-kb windows were low for both populations, but slightly higher for 33 wolves than for 67 domestic dogs (0.225 vs. 0.174; Fig. 3A). Limiting the dog panel to 42 Chinese village dogs yielded the same result. Over the whole chromosome, the result was sensitive to the metric, with raw-rate correlations being nearly tied but slightly favoring village dogs (0.552 vs. 0.534), and log-rate correlation favoring wolves (0.376 vs. 0.312). In map comparisons, *r* refers to Pearson correlation with the pedigree map; marker-pair dosage *r*^2^ is used only for the LD-decay diagnostic in Fig. 3B. We then expanded the comparison across chromosomes 1–5 to test whether the chromosome-1 result was part of a general trend. Correlations at 100 kb were 0.384 for wolves vs. 0.329 for village dogs, yielding a paired chromosome-bootstrap difference of 0.056 (95% interval 0.012–0.091). The corresponding log-rate correlations were 0.318 and 0.271, with a difference of 0.047 (0.028–0.065). The result for chromosome 1 was nearly tied but slightly favored dogs, whereas chromosomes 2–5 favored wolves. Thus, our comparison across chromosomes suggests an average advantage for wolves (SI Appendix, Fig. S4B). We next asked whether such an advantage persisted after controlling for multiple factors, including sample size, callability, marker density, and allele-frequency spectrum. We performed chromosome-1 map recovery using 20 matched panels of 33 wolves and 33 village dogs, matched for the total number of completely called variants within each 100-kb/0.05-MAF stratum. The median raw-rate correlation was 0.551 for wolves vs. 0.475 for village dogs, corresponding to a paired difference of 0.077 (joint panel-and-block 95% interval 0.019–0.147). Sensitivity analysis with increasing numbers of missing calls yielded the same ordering but a smaller raw-rate difference (SI Appendix, Fig. S4A). As an independent biological confirmation, we related chromosome-1 rates to GC content and transcription-start-site density within 100-kb windows, with joint adjustment for these features and SNP density. The standardized GC-content coefficient was positive for the village-dog LD map (0.303; 5-Mb block-bootstrap 95% interval 0.130–0.477) but uncertain for the wolf LD map (0.185; −0.073–0.398). Neither of the two LD maps retained TSS density as an independent variable after adjustment. Altogether, our results support a modest, metric-dependent advantage of the less bottlenecked wolf population for map recovery, without implying biological accuracy of the wolf map in every respect (SI Appendix, Fig. S4C).

**Figure 3.**
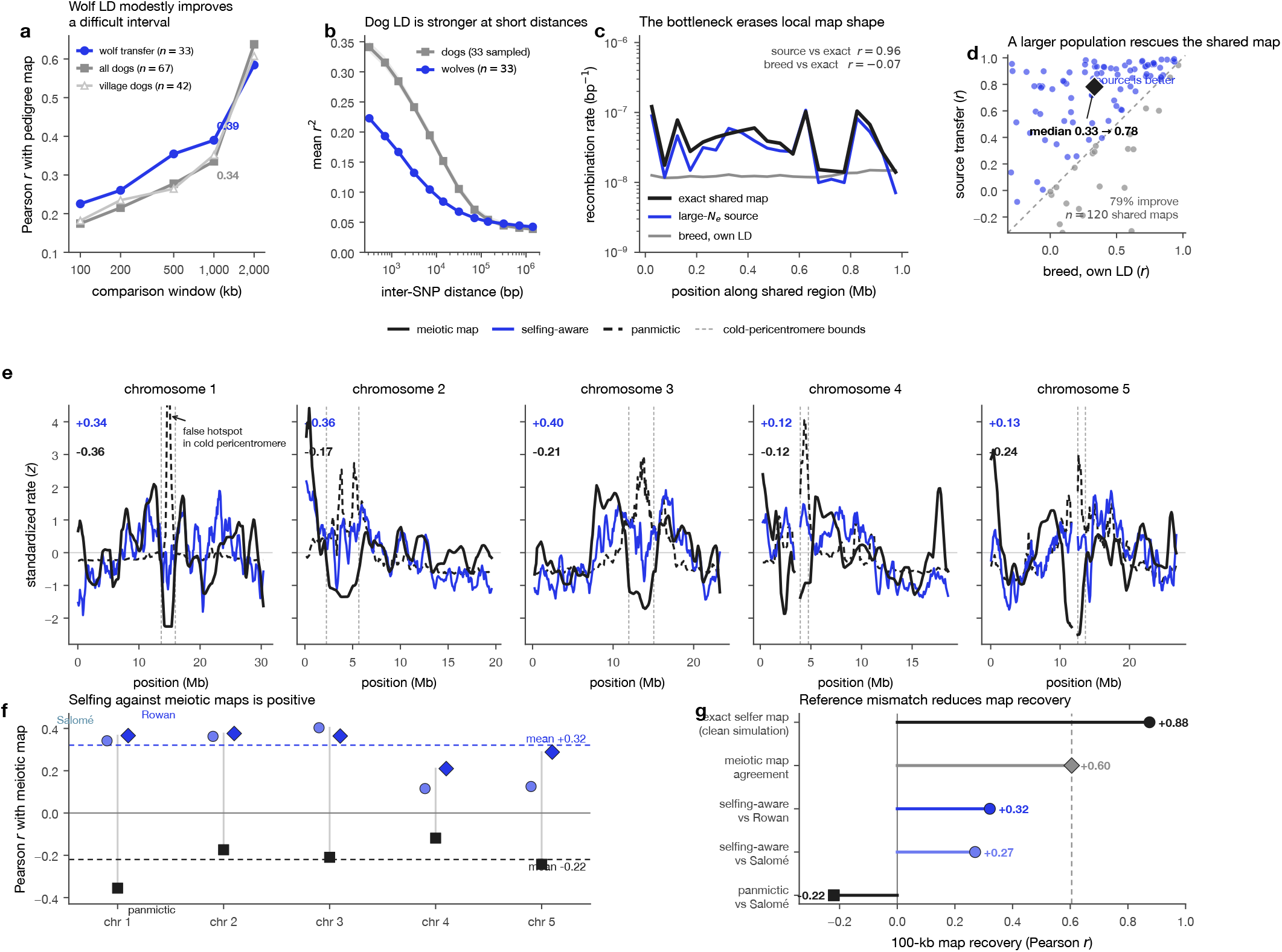
Biological boundary cases for LD-based transfer. (a) Pearson correlation with the Campbell dog pedigree map across reporting resolutions for chromosome-1 estimates from wolves, all domestic dogs, and Chinese village dogs. (b) Dosage *r*^2^ decay in 33 wolves and 20 sample-matched panels of 33 village dogs; shading shows the 2.5th–97.5th percentiles across dog panels. (c) Paired simulation with one large and one bottlenecked population using the same recombination map. (d) Pearson correlation with the underlying log-rate map across 120 pairs of simulations; the diamond shows the bivariate median. (e) Selfing-adjusted and random-mating population maps for all five *A. thaliana* chromosomes; dashed lines indicate pericentromeric regions. (f) Correlations with Salomé and Rowan meiotic maps on a per-chromosome basis. (g) Simulated recovery, map agreement, and empirical correlation.

In 120 paired simulations using the same recombination map, the bottleneck eliminated local peaks and valleys recoverable from the larger population (Fig. 3C; SI Appendix, Fig. S5A). At 100 kb, the median correlation with the true log-rate map was 0.33 when using the bottlenecked population and 0.78 when using the larger population (Fig. 3D). 95 out of 120 regions improved, with a paired median improvement of 0.22 (bootstrap 95% interval 0.16–0.39). Improvement from the bottlenecked population was positively associated with its current effective population size (Spearman *r*_*s*_ = −0.55, *P* = 1.1 × 10^−10^), whereas improvement from the larger population depended on the depth of the bottleneck (*r*_*s*_ = − 0.50, *P* = 9.0 × 10^−9^). Hence, simulations showed a mechanism for map shape recovery from a less bottlenecked related population, while empirical analysis revealed a small and metric-dependent difference between canids rather than the cause of this difference.

### Mating system and population structure require explicit modeling

As selfing modifies ancestry and the effective recombination rate, we considered it as a model limit warranted of explicit modeling. We examined 156 haplotypes from southern Sweden in the *A. thaliana* 1001 Genome dataset (30). When compared to the Salomé map, `fastrho` estimates accounting for selfing were positively correlated on all five chromosomes (*r* = 0.116–0.403; mean 0.270), while `pyrho` estimates using a random-mating assumption were negatively correlated on all five chromosomes (*r* = −0.355– −0.119; mean −0.220). The random-mating model generated additional peaks in regions with low recombination rates, while the selfing-aware model corrected for these peaks (Fig. 3E; SI Appendix, Fig. S5B).

On chromosome 1, replacing the population history in the random-mating `pyrho` lookup table with a constant population size, a two-epoch expansion, or a three-epoch bottleneck did not improve correlations of any kind, leaving them around −0.35 (SI Appendix, Empirical tests of demographic and mating-system effects). Demographic history alone therefore did not reproduce the effect of selfing. Across all five chromosomes, the selfing-aware map was positively correlated with the independent high-density Rowan map (*r* = 0.321; Fig. 3F) (31). Salomé and Rowan meiotic maps were consistent at *r* = 0.605, whereas a clean selfing simulation resulted in the recovery of the exact map used to generate the data at *r* = 0.875 (Fig. 3G).

We then tested the fixed selfing model against an independent linkage map in two non-model *Arabis* species (32). The reference contained 2,082 markers from 742 *F*_2_ offspring of reciprocal interspecific crosses (21). At 2 Mb, correlations with this shared cross map were *r*_*s*_ = 0.330 for 25 *A. sagittata* accessions (chromosome-bootstrap 95% interval 0.083–0.521; *P* = 0.0012), 0.158 for 12 *A. nemorensis* accessions (−0.204– 0.528; *P* = 0.189), and 0.305 for the chromosome-normalized geometric mean of both species ( −0.017–0.600; *P* = 0.0306; SI Appendix, Fig. S8). Excluding chromosomes 4 and 7, which show segregation distortion in the cross, increased the respective *A. sagittata*, combined, and *A. nemorensis* correlations to 0.443, 0.425, and 0.236.

A small-sample specialized model increased the *A. sagittata* correlation to *r*_*s*_ = 0.519 but did not improve *A. nemorensis* (*r*_*s*_ = −0.021). Twelve geographically distributed subsamples of 12 *A. sagittata* accessions all remained positively correlated with the cross map (median *r*_*s*_ = 0.592; range 0.515–0.633), whereas leave-one-out analysis of *A. nemorensis* remained near zero or negative (median *r*_*s*_ = −0.037; range −0.186 to −0.023). Changing the rate of selfing did not account for this discrepancy either. We therefore trained a second model on simulations that covered random mating, weak island model structure, both sampling designs used empirically, migration, split history, population size, and selfing. The model was trained based on simulation performance and chosen before the cross map was re-scored. For 2 Mb windows, the structured model gave correlations of 0.270 for *A. nemorensis* (chromosome-bootstrap 95% confidence interval −0.034 to 0.545; spatial-shift *P* = 0.046), 0.657 for *A. sagittata*, and 0.536 for their combined maps. Omitting the *A. nemorensis* accession used as an *F*_2_ parent raised the correlation for the *A. nemorensis* map to 0.354 (0.089; *P* = 0.0099). These post hoc follow-ups indicate that population structure drives experimental map correlation.

### A large-*N*_*e*_ specialized model scales inference to Ag3

The previous studies of LD and crossing experiments have shown the high-level resolution of recombination in mosquitoes, but they lack population-specific Ag3 population panel resolution to tens-of-kilobase resolution(33–37).

Therefore, we utilized the fixed large-*N*_*e*_ sampling method for phased Ag3.0 haplotypes from 13 pre-defined populations: six *Anopheles gambiae*, four *Anopheles coluzzii*, and three *Anopheles arabiensis* populations. We settled on analyzing 40 diploid mosquitoes for each population, having the same set of mosquitoes for 2R, 2L, 3R, 3L, and X chromosome arms, with rates summarized in nonoverlapping 50-kb windows (22).

If interaction between the inversion arrangements results in LD over a long distance, the predicted cold 2La block must be more pronounced in the populations that have both of the arrangements. The suppression depth was determined by the median ratio of recombination rate in the defined 20.524–42.166-Mb breakpoints in relation to the rest of 2L. Across 13 populations, suppression depth was positively but imprecisely associated with expected heterokaryotype frequency: Pearson *r* = 0.510, *P* = 0.075; Spearman *r*_*s*_ = 0.379, *P* = 0.201 (Fig. 4B,C). The three *Anopheles arabiensis* populations were nearly fixed for one arrangement and therefore contributed little variation at low expected heterokaryotype frequency.

**Figure 4.**
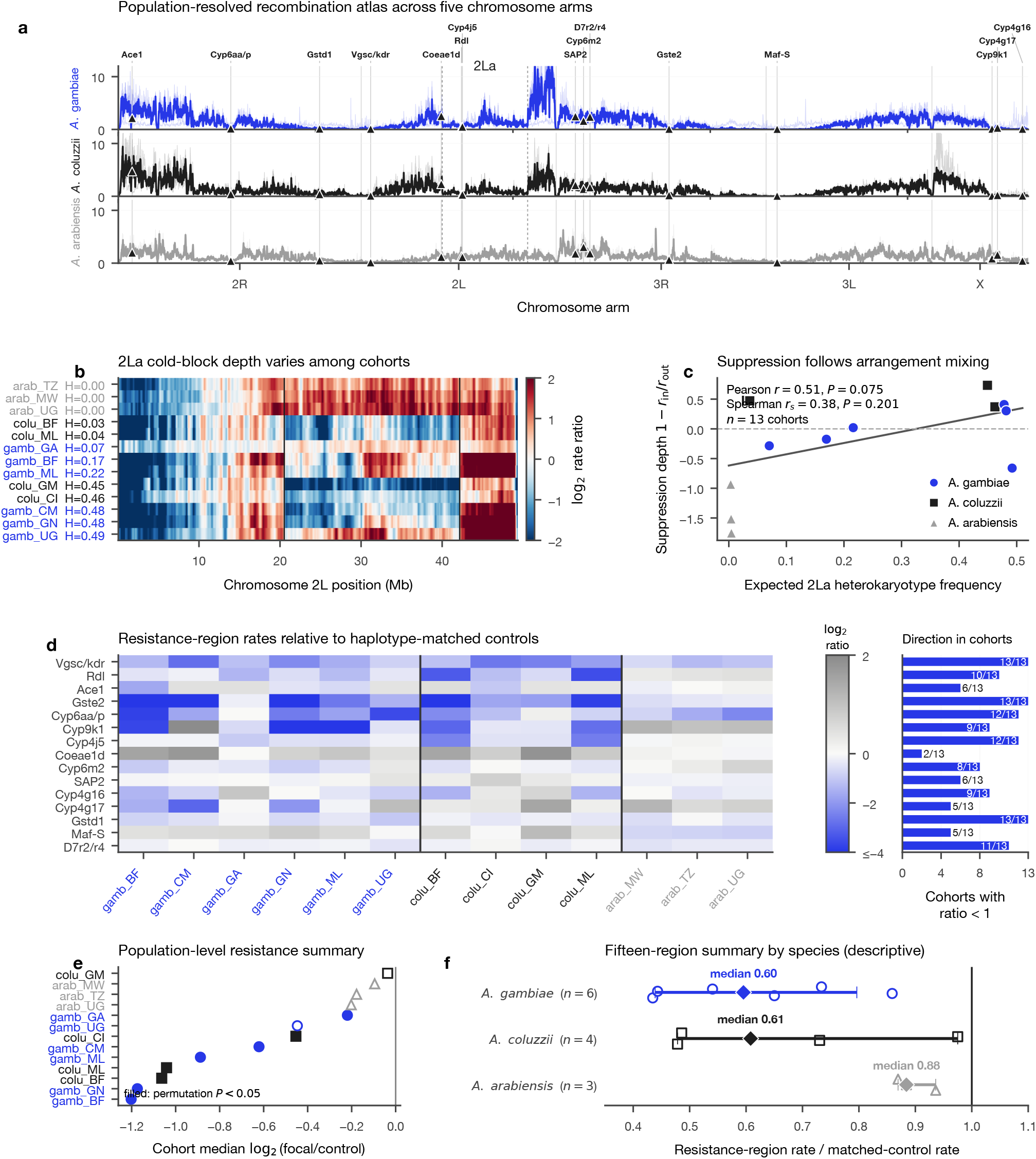
Recombination landscape and resistance region analysis of Ag3. (a) Recombination rates of 50-kb window sizes in 13 populations on five different chromosome arms. The thin lines are the population maps, while the thick lines are species medians and triangles indicate 15 literature-defined resistance regions. The normalization is done independently for each arm in each population. (b) Log_2_ rates of 2L compared with the median of 2L (but not 2La); rows have been smoothed for clarity. (c) Suppression depth versus heterokaryotype frequency in 2La (d) Log_2_ rate in 15 resistance regions and 13 populations matched by chromosome arm, position, SNP density, nucleotide diversity, and H12; numbers at the margin refer to the number of populations with ratio less than one. (e) Per-population geometric means of ratio; filled symbols show nominal within-population permutation *P* < 0.05. (f) Descriptive statistics for the same 15 resistance region statistic. Diamond indicates median and error bars indicate 95% percentile interval from 4,000 population-bootstrap samples.

### Ag3 pedigrees support fastrho’s LD based inference

The LD-inferred atlas was compared against an independent crossover map based on 15 Ag3 colony pedigrees (22). The two maps use independent datasets: wild mosquitoes were used to infer the atlas, while direct observations of transmissions in the crosses were used to generate the crossover map. The main analysis used five held-out crosses and autosomal 5-Mb windows; all 15 crosses and 2-Mb windows were used for sensitivity analyses.

Crossovers were called independently of the inferred atlas, its validation confirmed that the true crossover location was found within the reported breakpoint interval in 99.11% of simulation trials, and 3,959 of 4,000 introduced crossovers were successfully detected in the transmission backgrounds. The five held-out crosses had 97 crossover intervals spanning 170 parental transmissions. Using 5-Mb windows, the correlation between the crossover map from the five held-out crosses and the 10-population *Anopheles gambiae*/*Anopheles coluzzii* atlas reached *r*_*s*_ = 0.470, while using all 15 crosses gave *r*_*s*_ = 0.692 (Fig. 5).

**Figure 5.**
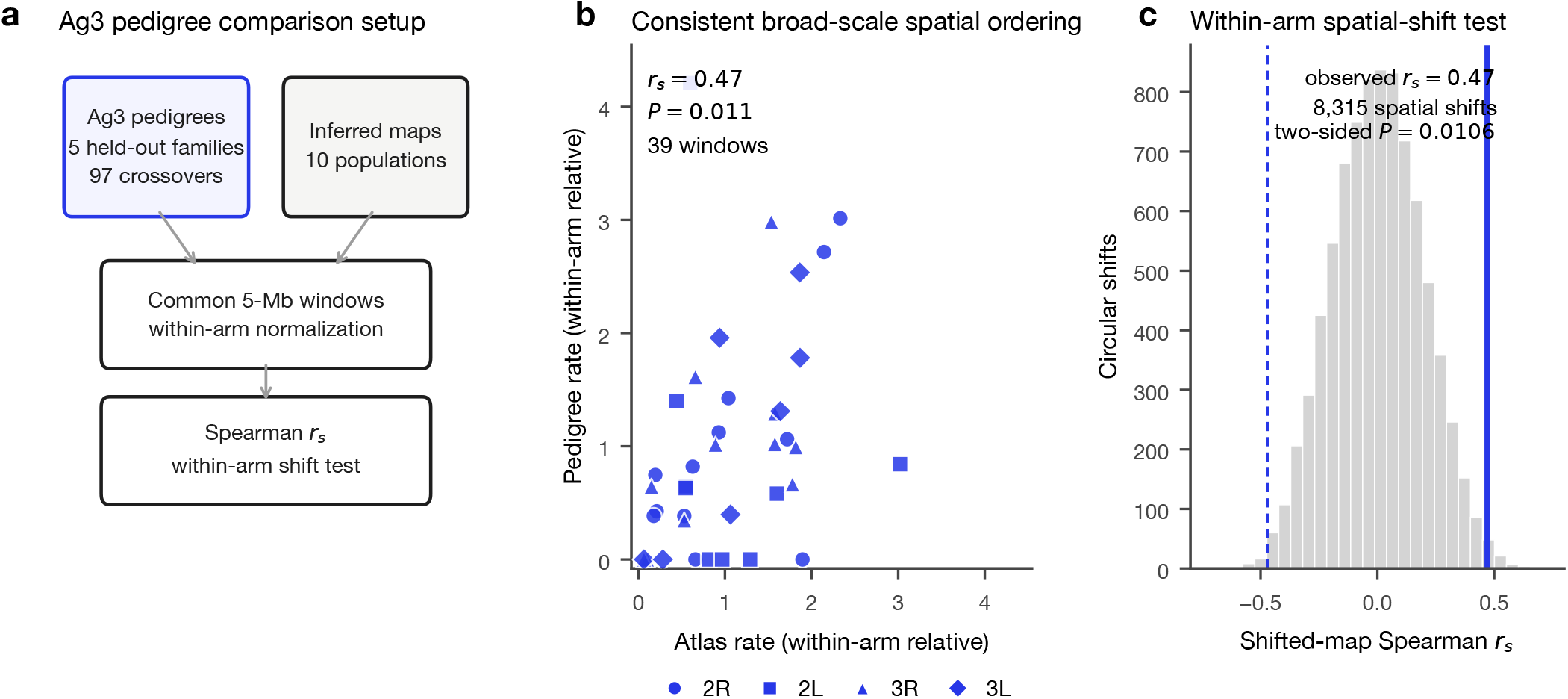
Broad-scale comparison with independent Ag3 pedigrees. (a) Crossover rates from five held-out families (97 crossover intervals across 170 parental transmissions) compared with the 10-population *Anopheles gambiae*/*Anopheles coluzzii* atlas in matched within-arm 5-Mb windows. Atlas estimates were unavailable during site filtering, phasing, crossover calling, and caller validation. (b) Chromosome-arm-normalized rates in supported autosomal windows. Marker shapes indicate chromosome arms. (c) Correlations at 5- and 2-Mb resolution using either the five held-out families or all 15 families.

### Resistance loci reveal selection-associated LD-map structure

We tested 15 literature-defined insecticide-resistance regions against up to 75 controls per locus and population (23). Controls were on the same chromosome arm, at least 0.50 Mb from every focal region, and matched for relative chromosomal position, SNP density, nucleotide diversity, and H12. The union of all focal regions was excluded from every control pool, and the population statistic was the geometric mean of the 15 focal-to-control ratios.

Across all 13 populations, the median population-level focal-to-control ratio was 0.730, with nominal within-population permutation *P* < 0.05 in eight populations (Fig. 4D–F). *Vgsc/kdr, Gste2*, and *Gstd1* had ratios below one in all 13 populations, while *Cyp6aa/Cyp6p* and *Cyp4j5* did so in 12. Descriptively, the median was 0.595 across the ten *Anopheles gambiae* and *Anopheles coluzzii* populations and 0.884 across the three *Anopheles arabiensis* populations.

### Arrangement stratification resolves the redpoll cold block

The chromosome-1 inversion occurs in two orientations, arrangements A and B. Principal-component analysis of the inversion interval separated 72 redpoll genomes into 37 A/A homokaryotypes, seven inferred A/B heterokaryotypes, and 28 B/B homokaryotypes (Fig. 6A). Because pooling arrangements can generate long-range LD that resembles reduced recombination, we inferred maps both from all 72 birds and separately from the two homokaryotype groups (Fig. 6B). Seven heterokaryotypes were insufficient for a separate map, so this analysis tests the effect of arrangement mixing rather than crossover suppression within heterokaryotypes.

**Figure 6.**
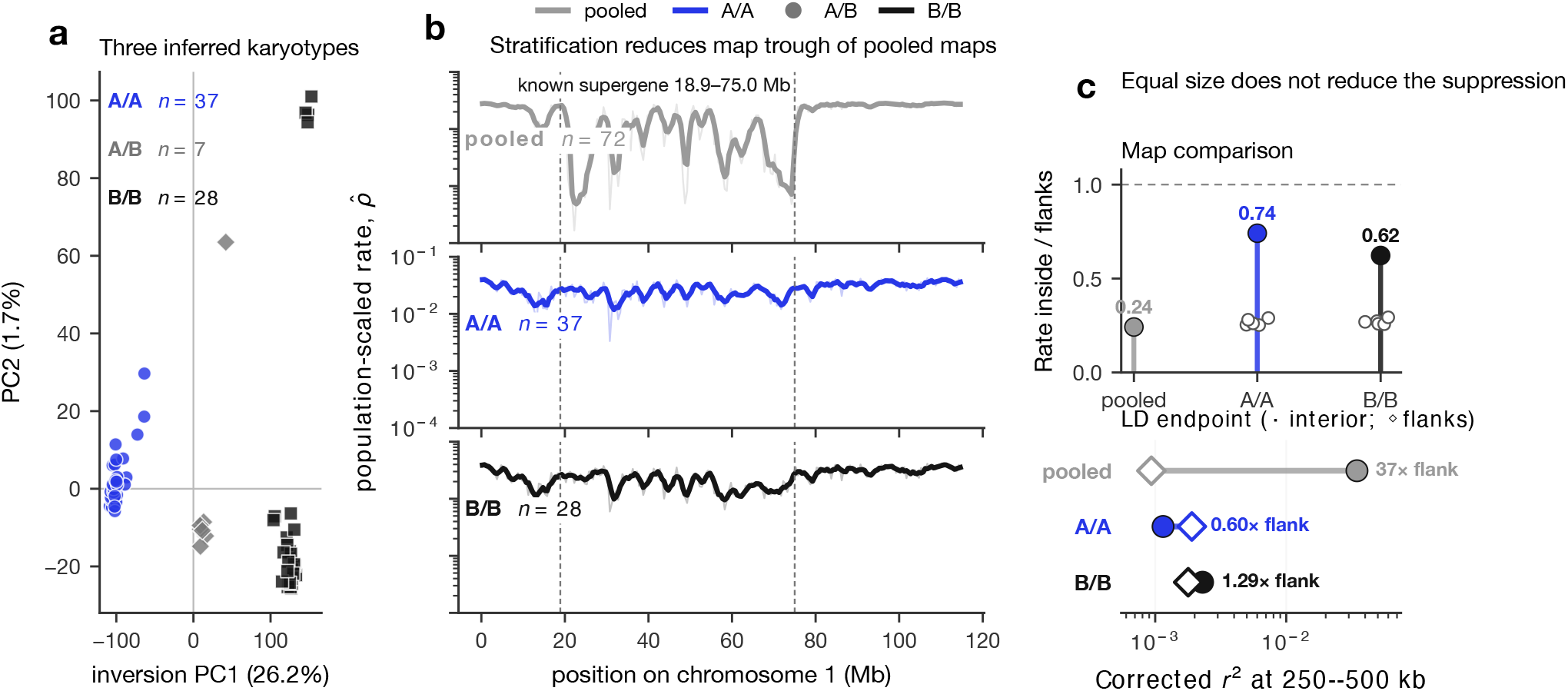
Arrangement mixing creates an LD-derived cold block in the redpoll supergene. (a) Principal-component analysis of the inversion interval separates 72 genomes into 37 A/A homokaryotypes, seven inferred A/B heterokaryotypes, and 28 B/B homokaryotypes. (b) Recombination-rate landscapes estimated from different groups of karyotypes. Thin lines show 500-kb inferences and thick lines show smoothed 2-Mb. The inversion is marked between 18.9–75.0-Mb . (c) Top, median inferred rate inside the inversion relative to the flanks; open symbols show five mixed-arrangement subsamples matched to each homokaryotype sample size. dosage *r*^2^ for variant pairs 250–500 kb apart inside the inversion (filled circles) and in the flanks (open diamonds).

The median inferred recombination rate in the inversion with respect to the flanks was 0.242 in the pooled sample but rose to 0.741 in A/A and 0.623 in B/B homokaryotypes (Fig. 6C). Mixed arrangements in subsamples equivalent in size to the smaller groups did not rise above 0.27, eliminating sample size alone as an explanation. An independently calculated genotype signal based on dosages for variants 250–500 kb apart showed corrected dosage *r*^2^ values within the inversion of 0.0346 in the pooled sample, 0.00115 in A/A homokaryotypes, and 0.00230 in B/B homokaryotypes. With respect to the flanks, long-range linkage disequilibrium was approximately 37 times higher in the pooled inversion but showed almost no enrichment in either homokaryotype. Thus, the lower level of recombination detected in the pooled sample is mostly explained by the composition of the sample rather than by a low level of crossing-over within either arrangement.

## Discussion

`fastrho` follows the simulation-based inference paradigm by simulating a landscape of possible evolutionary relationships using parameterized generative processes centered on a broad prior, here with the aim of fine-scale recombination mapping. Our estimator is designed to be transferable across datasets, accounting for variable LD patterns, which, as we have shown across many examples throughout this study, are shaped by the underlying biology. We therefore trained a broad base model on outbred populations and reused it across many demographic scenarios, including many species. In the same vein, we limited complexity by dividing evolutionary scenarios into common cases covered by the general model and cases requiring specialized models, such as selfing, as in *A. thaliana*; large population sizes, as relevant for mosquitoes; or mixed processes, such as selfing in combination with structure in the *Arabis* case. We therefore provide a general model alongside tools for quick and easy transfer to specific challenges.

Each real data application presented throughout this work represents a separate angle for LD and, by extension, recombination variation. Our cross-species analysis, for instance, demonstrated the general transferability on a species level. Here, the main confounders were variation in rates, that is mutation or recombination rates, or demographic fluctuations, those being more straightforward to integrate since they can be effortlessly simulated. More nuanced processes were likewise included, that is, more extreme bottlenecks, testing the boundaries of the amortization strategies while also showing the data loss based on biological and, frankly, theoretical limits, as shown here in the case of canids. The impact of severe population structure as another boundary was studied in *Arabis*, additionally layering complexity onto LD, particularly in combination with selfing, which reduces the effective number of recombination breakpoints. Then, our major contribution being the Ag3 recombination atlas, showing not only the inference scale, but also yet again the flexibility and benefits of amortized neural method development by scaling to large-Ne organisms. This result in particular has been independently verified by an available number of experimental crosses, showing correlation at a coarser level due to the available number of crosses. And lastly, we ended by showing the effect genomic structural variation and the pooling of different underlying structures have on local genomic correlation, presented through visualizing the redpoll supergene regions. All these individual applications provide an excerpt of variation that can be studied when looking through the prism of recombination.

In the Ag3 dataset, we studied two types of recombination rate heterogeneity in *Anopheles gambiae*, the first one driven resistance-associated selection and second one by structural variation, respectively. Both are known for increased LD and locally decreased recombination leading to generation of longer and more correlated haplotypes in comparison to the background (38, 39). On average across 13 populations and all five chromosomal arms, 15 insecticide-resistance regions showed the recombination rate shift to lower values compared to the haplotype-matched control regions. The magnitude was varied for all loci and populations, implying that the evolution of resistance affects the recombination landscape differently, in other words, the impact on the recombination landscape is allele, region and evolutionary history dependent. SNP density, nucleotide diversity and H12 levels were matched between test and control regions in order to minimize the effect of selection. The most straightforward example of the effect of structural variation and population composition on the recombination landscape is the case of 2La inversion (40). The LD and recombination landscapes associated with it significantly depend on the karyotype mixture in particular populations. Pooling individuals with different karyotypes results in generation of extended haplotypes and long-range LD and hence can result in recombination cold region although patterns within individual karyotypes are weaker. It implies that 2La is not an inherent feature of the inversion but rather a result of interaction of the inhibited exchange among different genomic arrangements and the frequencies of those in a population. In conclusion, resistance evolution and structural variation are key factors in the shaping haplotype variation. The remaining question is whether selection leads to the generation of LD alone and, consequently, to inferred decreased recombination rate or whether regions with low recombination rate provide favorable allelic combinations and hence help the process of selection?

The redpoll supergene provides an independent observation of this arrangement interpretation. A ∼55-Mb chromosome 1 inversion linked to plumage coloring and bill morphology, providing a mechanism for maintaining coordinated phenotypic variation despite ongoing gene flow among ecotypes (24). In our analysis, the broad recombination cold block was substantially stronger when alternative arrangements were pooled than within either homokaryotype. The same contrast in raw dosage LD show that this pattern was not solely an artefact of recombination-rate inference and pooling different arrangements increased LD correlations and thereby amplified effective reduction in recombination. Since only seven heterokaryotypic genomes existed, preventing any direct measure of crossover suppression, the pooling analysis clearly shows that mixture of arrangements can create a wide cold block from LD. Like 2La, this is probably due not only to current crossover frequency but also to the frequencies of the karyotypes themselves.

Canids and selfers expose two different limitations, which both lower effective model transfer through limiting available information between sites. Canids, due to their bottleneck, show a reduced number of polymorphisms and a scale shift in LD, which is why we tested whether information could be recovered from a closely related species such as wolves. In fact, correlations of our inferences with the dog pedigree map were higher in wolves throughout multiple chromosomes. Selfing underlies a fundamentally different mechanism, yet also leads to the reduction of effective recombination observability. Here, meiosis still leads to crossovers, but due to increased homozygosity, sites appear more correlated, and added population structure further increases covariances between the underlying local genealogies. Therefore, accounting for this particular mating system in *Arabidopsis* was vital for the respective landscape recovery, and accounting for population structure in *Arabis* was likewise essential to increase experimental map correlation (32). Therefore, it becomes essential not to disregard specialized models, which are fine-tuned to the particular biology at hand.

The model estimates the population-scaled rate *ρ* = 4*N*_*e*_*r*, and not the absolute rate *r*. The use of *N*_*e*_ in this way is very different from conditioning a lookup table or target-specific estimation algorithm on a single demographic history. `fastrho` does not require *N*_*e*_ or a demographic history during inference; instead, it has generalized based on a wide range of simulated histories and translates the LD into a value for *ρ*. Any provided *N*_*e*_ is used only later to calculate *r* = *ρ*/(4*N*_*e*_). Thus, using *N*_*e*_ could be replaced with *cN*_*e*_, resulting in a rescaling of all estimates, but without leading to changes in the estimated overall rate distribution. By contrast, with demographic lookup tables or target-specific training, changing the demographic history can change the mapping from LD to *ρ*. This will reduce the sensitivity of inferring the map shape to the assumed demographic history, while the sensitivity of inferring absolute values is maintained with respect to both *N*_*e*_ and the mutation rate. The intervals reported do not incorporate any uncertainty associated with these values. Simulations for training included crossovers but not gene conversion; longer conversion tracts interfered with map shape inference and increased the estimated rate.

Within these limitations, `fastrho` creates a valuable population-genomic perspective to experimental recombination mapping. Crosses and pedigrees remain the gold standard due to their direct crossover observation, or for providing independent inference evidence, but our resuseable LD estimator extends map construction to populations and species for which such experiments are impractical. Maps produced by such a procedure can be used to prioritize regions for further investigation, as well as to conduct analyses among various populations, as long as absolute scaling is conditional upon *N*_*e*_ and that biological processes affecting LD are evaluated explicitly.

## Online Methods

All models and their training datasets, together with the composition and sources of the empirical datasets, are summarized in SI Appendix, Tables S5, S9, S1, and S2.

### Tokenization and model

Inputs were defined at per-variant resolution. Each biallelic segregating site *i* was associated with the interval [*p*_*i*_, *p*_*i*_+_1_) to its right and represented by 17 features:

- four geometry and frequency features: distances to the preceding and following SNPs, derived-allele frequency, and minor-allele frequency;
- six multiscale LD features: mean *r*^2^ and the number of SNP pairs contributing to each mean within cumulative radii of 5, 25, and 50 kb across both flanks;
- five adjacent-site features: *r*^2^ and the frequencies of the four possible allele pairs, *AB, Ab, aB*, and *ab*; and two local-polymorphism features: mean diversity and haplotype richness within a ± 8-SNP neighborhood.

Up to 200 neighboring SNPs were used in each direction for the LD calculations. To make the model work under the same conditions for all input types, we trained it with phased, unphased, and unpolarized input. For the latter two types, haplotypes were replaced by diploid genotype dosages *G* ∈ {0, 1, 2}. LD between loci *i* and *j* in the resulting representation was quantified by corr (*G*_*i*_, *G* _*j*_)^2^, whereas gametic covariance was obtained from the dosage covariance by *D*_*i j*_ = cov (*G*_*i*_, *G* _*j*_)/2 (41). We then obtained the composite feature *p* _*AB*_ = *p*_*i*_ *p* _*j*_+*D*_*i j*_, with the other three features derived from the marginals as *p* _*Ab*_ = *p*_*i*_ − *p* _*AB*_, *p*_*aB*_ = *p* _*j*_ − *p* _*AB*_, and *p*_*ab*_ = 1 − *p*_*i*_ − *p* _*j*_+*p* _*AB*_. This made it possible to use the same input layout for unphased data, although haplotype richness was substituted with its phase-invariant counterpart based on the number of unique diploid genotypes in the neighborhood. In the absence of an ancestral allele, any locus at which the counted allele frequency exceeded one half was included in the minor-allele encoding of the feature by transforming the dosage encoding *G* to 2 − *G*. For the base checkpoint, there were 17 features per token considered. In the joint-checkpoint, trained over phased, unphased dosage, and folded-unpolarized representations, one additional feature was included, namely the fraction of sites where the minor allele count was two or less. Two conditioning variables provided information on whether the input was dosage or whether the alleles had been folded. This information was provided via the conditioning vector rather than being concatenated with the token features. Both the neighborhood-mean and the *r*^2^ of the adjacent site were adjusted by subtracting the finite sample adjustment 1/*n* floor and then constraining the result to be greater than or equal to zero. As different numerical feature distributions arise from the phased, unphased, and unpolarized input types, the input was normalized using the mean and standard deviation calculated from its respective training set. During inference, the feature definitions and size of the input were compared against the expected input by the model and incompatible input was not processed.

The inputs were initially mapped to 256 channels, followed by mixing the local context using parallel convolutions of sizes 3, 7, and 15. Six encoder and four decoder layers were used in the model architecture; each of them was based on the bidirectional Mamba-2 block (42, 43). The input was separately scanned in the forward and reverse directions in the block, after which the representations were combined. Residual connections and feedforward layers 256 → 1024 → 256 (GELU) were incorporated into each block. The outputs of the encoders were then fed into the decoders in reverse order of depth from the deepest encoder representation to the shallowest one. The initial and final encoder blocks took as input the values of log_10_ *μ* and log_10_ *n*_hap_ representing the mutation rate and the number of sampled haplotypes, respectively. In addition, for the model trained using input variants, two additional features denoting the input being based on the dosage and alleles being folded according to the minor allele frequency were supplied to the model.

For each valid interval, the model predicted the mean and log variance of the standardized log *ρ*. During training, the main loss was a beta-weighted Gaussian negative log likelihood (44), to which squared-error losses for the region-level log *N*_*e*_ and mean log recombination rate were added, each with a weight of 0.1. Sequences longer than the 1,024-input context were divided into overlapping chunks with an overlap of 256 inputs. Predictions from overlapping chunks were averaged using positive Hann weights, which downweight chunk edges and give greatest weight to each chunk’s center. For this purpose, both the predicted means and second moments were combined, where *E*[*X*^2^]= *σ*^2^+*μ*^2^, so that disagreement between the predictions of different chunks also contributed to the final variance. After transforming the prediction back from its standardized form, the point estimate of the absolute recombination rate was calculated as 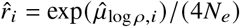. At inference, *N*_*e*_ was either supplied by the user or taken from the model’s auxiliary region-level *N*_*e*_ head; because mutation rate is an input to the model, either route remains conditional on the assumed mutation rate. The corresponding 95% limits were calculated as 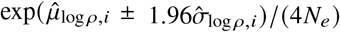. These limits were conditioned on a single value of *N*_*e*_, and therefore did not include uncertainty in *N*_*e*_ or mutation rate.

For unphased inputs, the four *p*-quantities are composite frequency features derived from dosages. The checkpoint was trained with this same feature definition. The model size along with their specified inputs and training priors used for the base and specialized versions are given in SI Appendix, Fig. S2 and Table S9.

### Simulations and benchmark scoring

The training and test regions were generated using `msprime` with explicit variable `RateMap` objects (25). For each simulated region, the exact recombination map used to generate the data was retained and later used as the reference for evaluation. The simulations included constant, autocorrelated, hotspot, species-specific, and published-map scenarios. The published profiles consisted of the deCODE and HapMap human maps, the Campbell dog map, the Comeron *Drosophila* map, and the Salomé *Arabidopsis* map (26–28, 45, 46).

All six `ReLERNN` models used phased input, the generating mutation rate through `--assumedMu`, `--maxSites 1750`, and the documented `--upperRhoThetaRatio` parameter, set prospectively to 3 for the five human-like scenarios and 8 for dog; no input-map percentile was used. Each model used 100,000 training, 10,000 validation, and 10,000 test simulations, 100 epochs, seed 1, and the generating demographic history.

For the window-based evaluation at the `ReLERNN` window scale, we generated 20 independent 10-Mb test regions of 20 phased haplotypes per scenario, each with one constant recombination rate fixed before simulation as the length-weighted mean of the corresponding source map. The trained models were reused without retraining. Exact truth, `fastrho`, and `pyrho` predictions were averaged within each complete `ReLERNN` output interval. Separate constant-demography `pyrho` and `ReLERNN` arms for bottleneck and expansion used identical test VCFs and windows. Fig. 2A reports the core `ReLERNN` predictions; the optional bias-corrected stage is reported in the SI Appendix.

For the staged timing comparison, we used the same 20 phased 10-Mb bottleneck regions as the window-based accuracy benchmark. All three methods were run sequentially in one Slurm allocation on one physical NVIDIA B200 node with 28 allocated CPU threads and 224 GB RAM. `fastrho` timing separated checkpoint loading from prediction; upstream simulation and pretraining were not repeated per dataset. `pyrho` timing separated construction of the matched-demography lookup table from inference of the 20 maps. `ReLERNN` timing separated simulation of the 100,000/10,000/10,000 training, validation, and test sets, 100-epoch training, and prediction.

For the window-based bottleneck and expansion controls, the primary `ReLERNN` model and `pyrho` lookup table used the generating history; a deliberately misspecified arm used a constant history. Each pair used identical test VCFs, truth maps, and complete `ReLERNN` output windows. Recovery of map variation was evaluated with Pearson and Spearman correlations. Absolute-rate error was summarized by the median predicted-to-true ratio, and interval coverage by whether the estimated interval contained the true recombination rate.

Only crossover events were included in the training data and in the main simulation experiments; gene conversion was not included in these simulations. For the separate gene-conversion test, we simulated 24 paired maps based on four gene-conversion initiation-rate ratios and three mean tract sizes. The trained model was kept fixed without retraining, while *N*_*e*_ was provided at prediction time, and the resulting predictions were evaluated at a resolution of 25 kb. The detailed simulation design and bootstrap procedure are provided in the SI Appendix. The linked-selection tests were performed separately using SLiM 5 (47).

### Canid bottleneck transfer

We built 120 pairs of canid populations sharing the same recombination map but differing in demographic history: one member had a large-population history and the other a recent bottleneck. This paired design isolated the effect of the bottleneck on recoverability of the simulated map.

For the empirical analysis, unphased genotypes from the Plassais panel and the Campbell domestic-dog pedigree map were used, both in reference to the CanFam3.1 genome assembly (28, 29). The displayed 40-Mb analysis involved 33 wolves and 67 domestic dogs. The Plassais panel and the domestic-dog pedigree map were each inferred with the same bottleneck-aware checkpoint, aligned together at the pedigree map level using 100-kb windows, and aggregated to coarser levels. The uncertainty in the displayed window was measured using resampling of eight adjacent 5-Mb windows. Dosage *r*^2^ was used independently to describe LD decay among variants with minor-allele frequency 0.05–0.95; 20 dog replicates were kept 33 individuals to match the wolf sample size.

In the expanded analysis, inference maps was done for 33 gray wolves and 42 Chinese village dogs on chromosomes 1 to 5. The correlations were calculated using windows of 100, 200, 500, and 1,000 kb pairs, and the chromosomes were resampled as paired chromosomes 10,000 times. In the chromosome-1 sensitivity analysis, we used 20 deterministic replicates used 33 village dogs, biallelic genotypes, and variants with a minimum MAF value of 0.05. For each 100-kb window and each 0.05 wide MAF bin, both populations were subsampled to the lower number of variants, followed by the inference using the frozen canid checkpoint.

For biological validation, CanFam3 chromosome-1 estimates were aligned to 100-kb GC summaries and unique transcription start sites from the UCSC Genome Browser. Standardized log rate was regressed jointly on GC, log(1+TSS count), and log(1+SNP) count for the Campbell, wolf-LD, and village-dog-LD maps. We obtained 95% intervals by resampling 5-Mb genomic blocks 10,000 times. We did not perform a PRDM9-motif analysis because PRDM9 is not functional in dogs, wolves, or coyotes (48, 49).

### *Arabidopsis* selfing

For the selfing analysis, we used 156 south-Swedish *A. thaliana* haplotype sequences and the Salomé/TAIR10 and Rowan meiotic recombination maps as references (30, 31, 46). Five chromosomes were analyzed using one selfing-aware model. The `pyrho` inference used one lookup table while varying the population-size history and retaining a random-mating assumption. Correlations were calculated in adjacent 100-kb windows.

### *Arabis* hybrid-cross benchmark

Population WGS panels were rebuilt using paired-end whole-genome sequencing data for 37 ingroup *Arabis* accessions (32). When species assignments provided in the ENA metadata contradicted the information in the original paper, we used labels in Supplementary Table S2 of the original paper. Reads were aligned to the chromosome-level reference genome of *A. nemorensis* GCA 976985625.1. Genotypes that had fewer than 5 or more than 60 reads were masked. Sites were filtered to complete, homozygous biallelic variants, and one allele was kept per locus for each predominantly selfing accession. This yielded separate population-WGS panels with 12 *A. nemorensis* and 25 *A. sagittata* genomes. For the initial test, both panels were analyzed with the fixed selfing model trained with 50–200 lines without any *Arabis*-specific retraining.

The reference map was assembled from 2,082 Restricted-site Associated DNA (RAD) markers detected in 742 *F*_2_ individuals from reciprocal interspecific crosses (21). All *F*_2_ individuals constituted a single interspecific linkage map and were not split into groups by parental species. Therefore, population-WGS maps were built separately for *A. nemorensis* and *A. sagittata* and were compared against the same linkage map, with coordinates based on the *A. nemorensis* assembly. Processing of the linkage map included collapsing duplicate positions, orienting chromosomes, and making genetic positions nondecreasing via isotonic regression. Within each chromosome, each map was divided by its mean because population-based estimates and the scale of a hybrid-cross map needed to be on the same scale. Our primary analysis used Spearman correlations in 2-Mb windows. We analyzed the results for each species separately and for the geometric mean of the two normalized inferred maps. In the controls, the analysis was repeated using 1- and 5-Mb windows, excluding distorted chromosomes and/or the *F*_2_ parents, excluding individual chromosomes, resampling chromosomes to estimate confidence intervals, and circularly shifting windows for the null test. The linkage map includes segregation distortion and was therefore treated as an independent empirical comparison rather than the species-wide ground truth.

In the first follow-up analysis, only the simulated panel-size range was changed from 50–200 to 8–50 lines. Five models were trained for 40 epochs each using 8,000 training and 800 validation regions. The models were selected based only on their performance in the simulations and were fixed before the empirical inference was performed. The final estimate was the average of their log-rate predictions.

The second follow-up employed 16,000 training, 1,600 validation, and 1,600 independent test regions. The simulations comprised a combination of 30% random mating, 30% weak island model, 20% the real *A. nemorensis* sample design, and 20% the real *A. sagittata* sample design. For these simulations, we varied the selfing rate, migration, split history, and number of individuals. Seven models were chosen based solely on their validation correlations with the simulations. Before scoring against the *F*_2_ map, the models had to satisfy predetermined criteria for both general and group-wise simulations in 100-kb windows; all seven satisfied these criteria. The same *Arabis* inference and scoring scheme was then conducted once. Both follow-up analyses were planned after inspecting the first empirical result and thus cannot be regarded as independent verification.

### Ag3 haplotypes, atlas, and 2La analysis

The mosquito atlas used the MalariaGEN Ag3.0 phased-haplotype release on AgamP4. We analyzed 13 predefined populations comprising six *Anopheles gambiae*, four *Anopheles coluzzii*, and three *Anopheles arabiensis* populations. Forty diploid mosquitoes were sampled without replacement from each population, and the same individuals were retained on all five chromosome arms. We selected biallelic segregating sites, and complete in the selected samples, and neighboring-interval estimates were aggregated into nonoverlapping 50-kb windows (22).

The 2La interval was 20.524–42.166 Mb on AgamP4. Within each population, the median positive rate inside the inversion was divided by the median positive rate across the remainder of 2L, and suppression depth was defined as one minus the ratio. Expected heterokaryotype frequency was estimated from all eligible Ag3 arrangement calls in each population, comprising 41–416 mosquitoes per population and 1,959 in total. Two-sided Pearson and Spearman tests related suppression depth to expected heterokaryotype frequency across the 13 populations (50–52).

### Ag3 pedigree crossover comparison

In the pedigree analysis, a total of 297 unique samples from 15 Ag3 crosses from the Ag3 colony were utilized; according to the publicly available metadata for the strains, the taxon is unassigned and hence specific identification to species level is not possible. In the first experiment, five Ag3 crosses which were not included in the development of the Ag3 site filter, 5 Mb windows on the autosomes, and crossovers less than 1 Mbp long were used. Two-state hidden Markov model estimated transmission of haplotypes from parents; phase block was divided by replicated marker disagreement; and any switch needed at least three markers called on either side. Atlas estimates were unavailable during site filtering, phasing, crossover calling, and caller validation. The rates were normalized within the chromosome arm and correlated using Spearman correlation, and sensitivity analysis was carried out using all 15 crosses and 2-Mb windows. Further information on caller validation is provided in the SI Appendix.

### Resistance-region controls

The 15 AgamP4 regions were selected from phenotype-associated markers and named, uniquely mappable insecticide-resistance genes (23). The adjacent genes within one window of 0.15 Mb were considered as one region; ambiguous families, inversions, microbial taxa, and regions found in *Anopheles funestus* were not included. The union of all focal regions was removed from every control pool. Up to 75 controls per region were selected on the same arm and at least 0.50 Mb from every focal region by matching relative chromosome position, log SNP density, nucleotide diversity, and H12. The population statistic was the exponent of the mean log focal-to-control ratio across the 15 regions. We used 5,000 within-population draws for permutation values and 4,000 deterministic population-bootstrap samples for the species intervals in Fig. 4F.

### Redpoll arrangement-specific maps and raw LD

Redpoll whole-genome genotypes were taken from the study of Funk et al. and its open-access database (24). In the analysis, we used 72 unphased diploid genomes on chromosome 1 that were genotyped relative to the brown-capped rosy-finch reference genome. The PCA performed using the genotypes in the known inversion interval allowed us to define two groups of homokaryotypes with 37 and 28 individuals as well as 7 heterokaryotypes. The recombination maps were constructed for the entire combined sample and separately for each group of homokaryotypes, after which the results were aggregated into 500-kb windows. The main statistic was the median of the positive recombination rate between 18.9 and 75.0 Mb normalized by the median of the positive recombination rate in the collinear regions. Here, the collinear regions were the flanking regions outside the inversion that retained the same marker order and orientation between arrangements. In order to take into account the reduced sample sizes in homokaryotypes, five random mixed-arrangement subsamples were created for each sample size. Reproducibility of the split map was analyzed using one random division of each group of homokaryotypes, and the concordance between two divisions was calculated separately for the inversion and collinear regions.

For the LD analysis without a genetic model, we merged the rows containing pseudo-haplotypes into diploid dosage genotypes. SNPs with counted-allele frequency outside 0.05– 0.95 were excluded, which is equivalent to excluding SNPs with minor-allele frequency below 0.05. In each distance bin, up to 20,000 SNP pairs were sampled independently from the inversion region and collinear flanks. Mean dosage *r*^2^ for these pairs was estimated, then the finite sample floor term 1/*n* was subtracted, and the values were bounded below by zero.

### Use of generative AI

Generative AI tools were used for editorial and computational assistance. All resulting content was reviewed and verified by the authors, who take full responsibility for the manuscript.

## Data availability

All empirical datasets are publicly available. Accession numbers, sample definitions, reference maps and source publications are listed in the SI Appendix.

## Code availability

Source code, documentation, checkpoints, analysis scripts and results are available at https://github.com/kevinkorfmann/fastrho. The repository includes reproduction workflows and an artifact registry under https://github.com/kevinkorfmann/fastrho/tree/main/reproduce.

## Acknowledgments

This study utilized the MalariaGEN Ag3.0 phased haplotypes and colony-cross resources. We thank the *Anopheles gambiae* 1000 Genomes Consortium, investigators and field teams of the contributing partner studies, the communities from which mosquito samples were collected, and the MalariaGEN data production and coordination teams (22, 53).

We thank Iain Mathieson for detailed feedback on the framing and presentation of the manuscript, and Martin Donnelly for discussion of and permission to include the Ag3 analysis.

We thank Andrew Kern for feedback on the ReLERNN application.

We also thank the providers of the redpoll and comparative genomic datasets used in this study.

## Author contributions

K.K. conceived the study, developed the software, performed the analyses and drafted the manuscript. N.R. contributed the *Arabis* benchmark, biological interpretation and manuscript revision. S.M. supervised and coordinated the research and contributed to interpretation and manuscript revision. All authors approved the final manuscript.

