## Supporting Information for "A reusable neural approach to recombination mapping for model and non-model species"

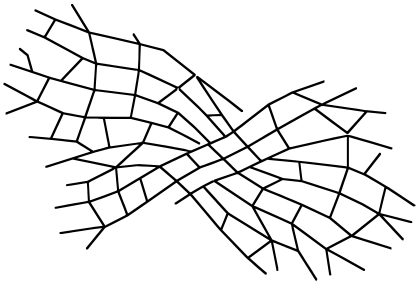

Kevin Korfmann[1,*], Neda Rahnamae[2], and Sara Mathieson[1]
[1]Department of Biology, University of Pennsylvania, Philadelphia, PA 19104, USA
[2]Institute of Plant Ecology and Evolution, Heinrich Heine University Düsseldorf, 40225 Düsseldorf, Germany

#### SI Overview

This Supporting Information contains method description and supplementary analyses for model evaluation, comparison across various species, canids, selfing, the Ag3 recombination atlas and pedigrees, insecticide-resistance regions, redpoll chromosome architecture, and *Arabis*. Table S1 lists the external datasets.

##### Data sources and versions

The main empirical sources were MalariaGEN Ag3.0, released on 4 November 2021, and the redpoll panel of Funk et al. in the Sequence Read Archive and Dryad. The main simulation tests used `msprime` 1.3.3, while the gene-conversion stress test used version 1.3.4. Table S1 records the other empirical and simulation sources and their versions. (1–4)

**Table S1:** Datasets and simulation resources used in the study. Versions and stable access routes are reported for each source.

| Source and version | Manuscript scope | Organism | Stable source | Citations |
|---|---|---|---|---|
| Ag3 Anopheles gambiae 1000 Genomes phase 3 haplotypes (3.0) | primary | Anopheles gambiae, Anopheles coluzzii, Anopheles arabiensis | MalariaGEN | (1, 2) |
| 1000 Genomes Project phase 3 genotypes (20130502 phase 3 release) | supporting | Homo sapiens | International Genome Sample Resource | (5) |
| Drosophila Genetic Reference Panel (DGRP2 freeze 2) | supporting | Drosophila melanogaster | NCBI Sequence Read Archive | (6, 7) |
| 1001 Genomes Arabidopsis accessions (2016 release) | supporting | Arabidopsis thaliana | 1001 Genomes | (8–10) |
| Plassais whole-genome canid panel (722 canids 2019 release) | supporting | Canis lupus familiaris, Canis lupus | NHGRI Dog Genome Project | (11) |
| Redpoll chromosome-1 supergene whole-genome panel (Funk et al. 2021 chromosome-1 panel, accessed 2026-07-20) | primary | Acanthis flammea | NCBI Sequence Read Archive and Dryad | (3) |
| stdpopsim species models and genetic-map catalog (0.3.0) | supporting | multiple | stdpopsim | (12) |
| msprime coalescent simulations (1.3.3 (primary benchmark) and 1.3.4 (gene-conversion stress test)) | primary | simulated populations | GitHub and PyPI | (4) |
| SLiM linked-selection simulations (5.0) | supporting | simulated populations | Messer Lab | (13) |

| Source and version | Manuscript scope | Organism | Stable source | Citations |
|---|---|---|---|---|
| Published human dog Drosophila and Arabidopsis genetic maps (versions listed in the cited sources) | supporting | Homo sapiens, Canis lupus familiaris, Drosophila melanogaster, Arabidopsis thaliana | stdpopsim catalog and primary publications | (9, 14–17) |
| Arabis nemorensis and Arabis sagittata $F_2$ linkage map and parental-species WGS panels (map commit 10c9092; curated 37-accession WGS panel; ENA accessed 2026-07-20) | supporting | Arabis nemorensis, Arabis sagittata | European Nucleotide Archive and GitHub | (18, 19) |
| Cross-species population-genotype comparison (ten species, accessed 2026-07-20) | supporting | mammals, insects, and plants | IGSR, DGRP, 1001 Genomes, EBI NextGen, ENA, and European Variation Archive | (5–8, 20–26) |

#### Simulation design and gene-conversion test

Training and test regions were generated with `msprime` using `RateMap` objects (4). Tests included constant-size, bottleneck, expansion, hotspot, and published-map scenarios based on the deCODE and HapMap human maps, the Campbell dog map, the Comeron *Drosophila* map, and the Salomé *Arabidopsis* map (9, 14–17). Species models used `stdpopsim` 0.3.0 (12, 27).

The training simulations modeled recombination but not gene conversion. The separate gene-conversion test used 24 paired 1-Mb regions, 12 with varying rates and 12 with discrete hotspots (28, 29). We simulated each region once without gene conversion and under 12 conditions combining initiation rates of 0.5, 1, 2, or 4 times the realized mean recombination rate with mean tract lengths of 100, 300, or 1,000 bp. All conditions used 20 haplotypes, $N_e$ = 10,000, mutation rate $1.5 \times 10^{-8}$ per bp, and mean recombination rate $10^{-8}$ per bp. We provided the fixed model with the known $N_e$ and evaluated predictions at 25-kb resolution. Pearson correlation measured map-shape recovery, and the paired ratio of mean inferred rates with and without gene conversion measured scale displacement. Intervals were obtained from 2,000 bootstrap resamples of the 24 regions (Fig. S1).

#### Phased input variables and prediction intervals

Let $s_1 < \cdots < s_S$ denote the positions of the chosen biallelic segregating sites, and let $H$ be an $n \times S$ haplotype matrix containing only 0s and 1s. The token $x_i$ points to site $i$, whereas the target of $x_i$ is the recombination rate per generation between sites $i$ and $i+1$. The last site does not have a valid interval to its right and, therefore, is not included in the resulting map.

The first four features encode positional and frequency information: $\log_{10}(s_{i+1} - s_i + 1)$, $\log_{10}(s_i - s_{i-1} + 1)$, derived-allele frequency $f_i$, and $\min(f_i, 1 - f_i)$. Missing preceding and succeeding gaps at a sequence edge are assigned a value of zero before application of the +1 transform. For each pair of sites $i$ and $j$, the tokenizer computes $D_{ij} = n^{-1} \sum_k H_{ki} H_{kj} - f_i f_j$ and $r_{ij}^2 = D_{ij}^2 / [f_i(1 - f_i) f_j(1 - f_j)]$. Then, for each focal site, it computes the average $r_{ij}^2$ for neighboring informative pairs inside cumulative radii of 5, 25, and 50 kb, and stores $\log(1 + N_{i,R})$ for the corresponding number of informative pairs. A pair within 5 kb is thus included in all three cumulative features. We limited the number of neighbors to 200 polymorphic sites to avoid featurization costs.

The exact adjacent two-locus state is preserved in five other channels: $r_{i,i+1}^2$ and the four haplotype frequencies $p_{AB}$, $p_{Ab}$, $p_{aB}$ and $p_{ab}$. The latter sum to one and are the normalized $2 \times 2$ haplotype-count table underlying two-locus composite-likelihood methods. The last two base channels are summaries of polymorphism in the ±8-SNP neighborhood. The per-site nucleotide diversity is $2c_i(n - c_i)/[n(n-1)]$, where $c_i = nf_i$, and the local channel is the centered rolling mean. Haplotype richness is the number of unique haplotype rows in the same neighborhood divided by $n$. The ordered 17-feature base token shown in main-text Fig. 1A results from concatenation of 4 geometry/frequency, 6 multi-radius LD, 5 adjacent-state, and 2 local-polymorphism channels.

The model trained jointly on phased, unphased, and unpolarized input representations uses 18 input variables per site. The extra feature is the local ratio of sites where the minor allele is observed at most twice in the sample. The model also corrects each mean $r^2$ and adjacent-site $r^2$ value for finite-sample noise by substituting $\max(r^2 - 1/n, 0)$ for it, where $n$ is the number of sampled haplotypes. This correction subtracts the expected small positive $r^2$ value from the sampling noise, setting any negative result to zero. The additional feature and LD correction were used during training and are required inputs for this checkpoint.

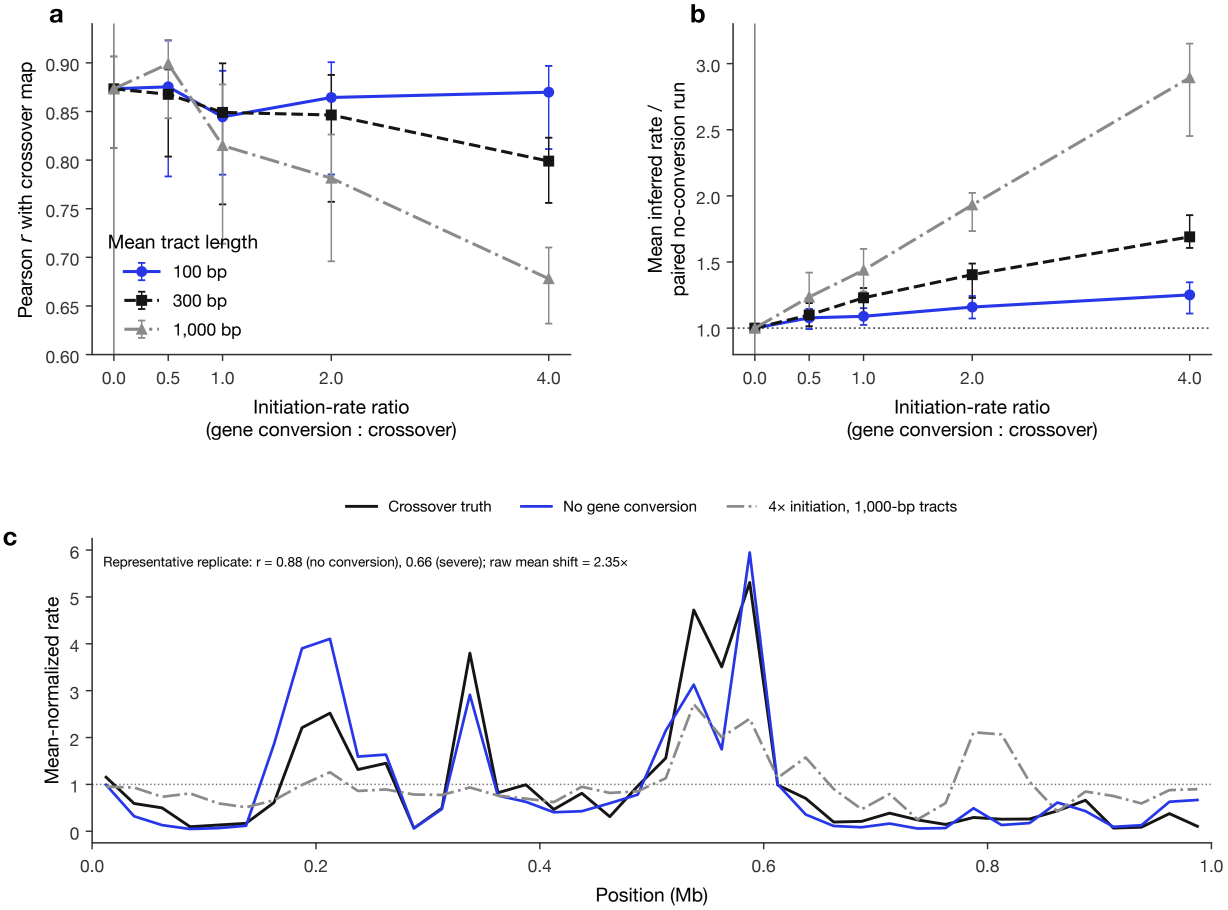


**Figure S1: Response of the fixed recombination-only model to gene conversion.** (a) Pearson correlation at 25-kb resolution with the exact recombination map across four conversion-to-recombination initiation-rate ratios and three mean tract lengths. (b) Median paired ratio of the region-mean inferred rate to its conversion-free counterpart. Error bars are 95% intervals from 2,000 bootstrap resamples of 24 recombination-map replicates. (c) Mean-normalized exact recombination map and predictions. Each curve is divided by its mean; the panel also reports the unnormalized mean shift and replicate-specific correlations.

#### Unphased and unpolarized input signals

For unphased data, the model estimates LD directly from diploid genotype dosages. The $n$ haplotypes are paired into $m$ diploid individuals ($n = 2m$) to produce the $m \times S$ genotype matrix $G$. The entries in $G$ are 0, 1, or 2, indicating the number of copies of the counted allele that an individual carries at a site. For the sites $i$ and $j$, we take the squared correlation of the genotype-dosage columns of sites $i$ and $j$, i.e., $\mathrm{corr}(G_i, G_j)^2$, rather than the $r^2$ of haplotypes. Under random mating, the covariance is estimated by $D_{ij} = \mathrm{cov}(G_i, G_j)/2$ (30). We then construct the composite frequency feature $p_{AB} = f_i f_j + D_{ij}$ and derive the other three features from the allele-frequency marginals: $p_{Ab} = f_i - p_{AB}$, $p_{aB} = f_j - p_{AB}$ and $p_{ab} = 1 - f_i - f_j + p_{AB}$. These are dosage-derived composite features. Local haplotype richness is replaced by the number of distinct diploid genotype patterns in the local neighborhood divided by $n$. All of these features are computed on genotype dosages, meaning that the resulting token sequence is independent of phasing information.

In cases there is no information on polarization, the model resorts to the use of the minor allele. Whenever the allele frequency exceeds one-half, dosage is recoded as $G_i \leftarrow 2 - G_i$. We leave pairwise $r^2$, informative-pair counts, nucleotide diversity, and physical distances unchanged.

Both mean and standard deviation of features were calculated using the training dataset and kept fixed during validation and inference phase. Each input representation was standardized independently while transformations for the recombination rate and $N_e$ were shared between them. Each checkpoint maintains its own feature definition and pre-processing procedure. And each checkpoint stores its definitions and preprocessing parameters and rejects incompatible inputs. Figure S2 compares the three input representations.

Accuracy decreased when phase and ancestral-allele information were removed. However, the model trained on all input forms

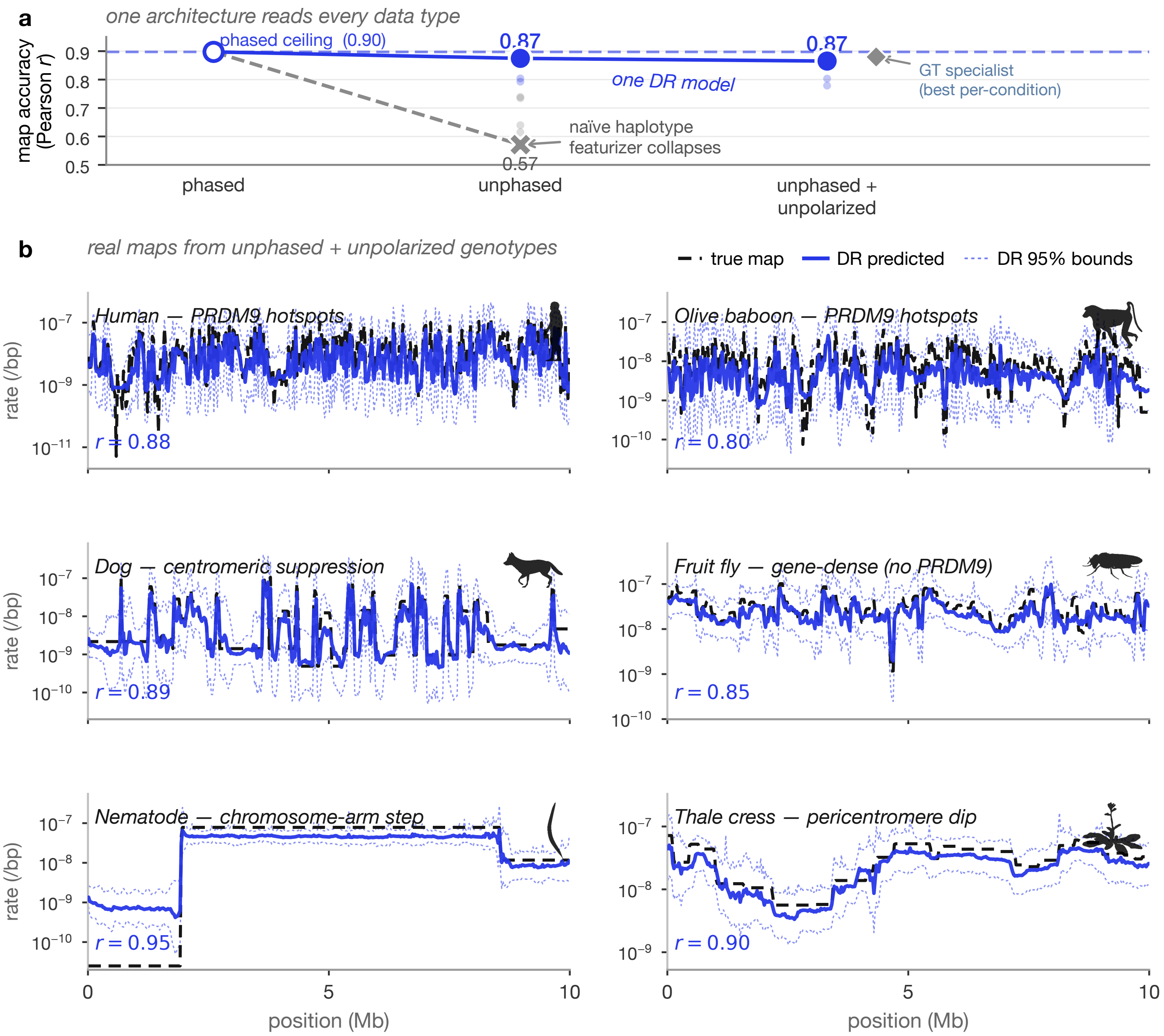


**Figure S2: Accuracy among forms of genotype input.** (a) Map accuracy for phased, unphased, and unphased-plus-unpolarized inputs, comparing the model trained across input forms with a phased-haplotype input used naively and with a separate model trained for each condition. Points display species, and larger symbols display their means. (b) Dashed reference maps, cobalt predictions from unphased, unpolarized data, and dotted conditional 95% prediction limits for representative human, canid, insect, nematode, and plant chromosomes.

still performed similarly to the phased model in the tested conditions (Fig. S2).

#### Bidirectional Mamba-2 encoder–decoder

The input to the `base-v1` checkpoint is a sequence of $L$ tokens, each defined by 17 standardized features. Each 17-feature token is linearly projected to 256 dimensions and then passed through a GELU activation. To let the model learn patterns spanning a variable set of distances, we use three one-dimensional convolutions having widths of 3, 7, and 15 tokens. The results of these three convolutions are then combined alongside the original 256-dimensional representation. The combined representation is projected back to 256 dimensions and passed through GELU activation and dropout. A learned positional embedding, that is, a trainable vector assigned to each position in a sequence, is then added so that the model can distinguish token positions in sequences of up to 1,024 tokens. The domain-randomized checkpoint has the same architecture but with 18 features per token instead of 17. Its conditioning vector also has two extra indicators that denote unphased genotype-dosage input and minor-allele folding.

The backbone is composed of six encoder blocks and four decoder blocks. Each block processes the layer-normalized tokens using two independent Mamba-2 state-space layers (31, 32), scanning them in both directions. The two 256-dimensional outputs are concatenated and projected back to 256 dimensions. The output is then passed through a dropout layer and added to the block input via a residual connection. After a second layer normalization, there is a 256 → 1024 → 256 GELU feed-forward network with dropout and another residual connection. This two-way design allows each token to use LD information from both sides while keeping the complexity linear in the sequence token length.

The output of each encoder layer is stored for use by the decoder, whose layers receive the projection vectors from the encoder layers in reverse order of depth, meaning the deepest encoder output is sent to the first decoder layer, and so on. After the first and last encoder blocks, feature-wise linear modulation is applied using a global conditioning vector containing $\log_{10} \mu$ and $\log_{10} n_{\text{hap}}$. For the domain-randomized model, this vector also includes indicators for genotype dosage and minor allele folding. The mutation rate may affect not only the mapping between genetic diversity and $N_e$ but also the model's internal sequence representation.

#### Training objective and model outputs

For the training target at interval $i$, we use the standardized log of the population-scaled recombination rate, $y_i = [\log(4N_e r_i) - m_\rho]/s_\rho$. The uncertainty head estimates a mean $\mu_i$ and a log variance $v_i = \log(\sigma_i^2)$. The interval loss is

$$\ell_{\beta,i} = \frac{1}{2}\,\mathrm{sg}[\exp(\beta v_i)]\left[v_i + (y_i - \mu_i)^2 \exp(-v_i)\right], \qquad \beta = 0.5,$$

where sg means stop-gradient, so the weighting term is treated as constant during backpropagation. This is the beta-weighted heteroscedastic Gaussian negative log likelihood (33). The full loss is the mean interval loss over valid intervals, plus two auxiliary terms: 0.1 times the squared error of the standardized region-level $\log N_e$ estimate and 0.1 times the squared difference between the predicted and true region-mean standardized log recombination rates.

The model has a recombination-rate distribution head and an auxiliary $N_e$ estimation head. At each token, we apply layer normalization to the rate head, followed by two separate two-layer GELU networks that predict the mean and log variance of the standardized log recombination rate. The $N_e$ head computes an average of the token representations for each sequence and feeds them to a dedicated multilayer perceptron to output one $N_e$ prediction for the entire sequence. The mutation rate input during inference influences the sequence representation as well as the auxiliary prediction, thus making the absolute scale conditional on this value. The uncertainty interval created by the rate head is a predictive interval conditional on $N_e$.

The estimated log population-scaled recombination rate after reversing the applied standardization is $\widehat{\log \rho_i} = \mu_i s_\rho + m_\rho$. Its standard deviation is $\hat{\sigma}_{\log \rho, i} = \sigma_i s_\rho$, which gives us the per-generation recombination-rate estimate $\hat{r}_i = \exp(\widehat{\log \rho_i})/(4N_e)$, using either a supplied $N_e$ or the auxiliary estimate. A nominal 95% predictive interval is $\exp(\widehat{\log \rho_i} \pm 1.96\hat{\sigma}_{\log \rho, i})/(4N_e)$. Given the known $N_e$ from a simulation, we assess interval coverage relative to the true recombination map used to generate the data.

#### Long-sequence inference

If a sequence is longer than the checkpoint context, it is split into overlapping chunks. The last chunk is not padded and can be shorter. During inference, overlapping predictions are merged using positively weighted Hann-interior coefficients, which give less weight to regions near the borders. Variance is merged using the weighted second moment, $E[X^2] = \sigma^2 + \mu^2$, so any differences between chunks will increase the estimated variance.

#### Benchmarking

The 25-kb `fastrho`–`pyrho` benchmark correlations are given in Table S6. The window-based results at the `ReLERNN` window scale are reported separately in Table S7. The ranges for the base and specialized `fastrho` simulation priors are defined in Table S9.

Pearson and Spearman correlations were computed to evaluate the recovery of variation of local maps at a given physical scale. We used the median estimated-to-true rate ratio as a summary of absolute rate error, while the area under the precision-recall curve was used to measure hotspot ranking. In each of the six benchmark situations, we simulated 2-Mb regions containing 20 haplotypes. For the constant, bottleneck, expansion, deCODE, and HapMap scenarios, the simulation parameters were as follows: $N_e$ = 10,000 and mutation rate $1.5 \times 10^{-8}$/bp/generation; for dog, $N_e$ = 13,000 and mutation rate $4 \times 10^{-9}$/bp/generation. Synthetic maps had a median crossover rate of $10^{-8}$/bp/generation and ranged from $10^{-10}$ to $2 \times 10^{-7}$/bp/generation.

Every `ReLERNN` model used `--maxSites 1750`, phased input, the scenario's generating mutation rate through `--assumedMu`, 100 epochs, and seed 1 (34). Training, validation, and test sets contained 100,000, 10,000, and 10,000 simulations, respectively. We used the documented `--upperRhoThetaRatio` parameter with prespecified bounds of 3 for the five human-like scenarios and 8 for dog; no input-map percentile was used. Each model used its generating demographic history. The first complete six-scenario suite was frozen before test scoring; correlations in the 10,000 held-out `ReLERNN` simulations were 0.957, 0.958, 0.906, 0.954, 0.953, and 0.948 for constant, bottleneck, expansion, deCODE, HapMap, and dog, respectively.

For the window-based test, we generated 20 independent 10-Mb regions of 20 phased haplotypes per scenario, each with one constant recombination rate fixed before simulation as the length-weighted mean of the corresponding source map. The frozen models were reused without retraining. Exact truth, `fastrho`, and `pyrho` predictions were averaged within each complete non-overlapping `ReLERNN` output interval, with terminal partial windows excluded for all three methods. There were no missing complete windows or nonfinite predictions. Nine finite `ReLERNN` predictions were exactly zero in the single deCODE region with the lowest generating rate ($1.02 \times 10^{-10}$/bp/generation) and were retained. Median complete-window widths were 0.74, 1.65, 2.85, 0.73, 0.74, and 2.28 Mb across the six scenarios.

Across the window-based suite, window-matched Pearson correlations ranged from 0.942 to 0.978 for `fastrho`, from 0.943 to 0.990 for `pyrho`, and from 0.847 to 0.946 for the core `ReLERNN` predictions (Table S7). Deliberately replacing the generating histories with a constant history changed `pyrho` from 0.943 to 0.892 and `ReLERNN` from 0.898 to 0.854 under the bottleneck; under expansion, `pyrho` changed from 0.961 to 0.967 and `ReLERNN` from 0.847 to 0.772. These arms used identical test VCFs and windows. The optional `ReLERNN_BSCORRECT` stage was run and independently rescored on the same windows; its `ReLERNN` correlations were 0.939, 0.883, 0.836, 0.933, 0.930, and 0.922 for constant, bottleneck, expansion, deCODE, HapMap, and dog, respectively. Main-text Fig. 2A uses the core `PREDICT` output because the documented workflow treats bias correction as optional, not because either stage was selected by its test correlation.

#### Canid bottlenecks

We ran simulations for 120 pairs of large and bottlenecked canid populations, all using the same recombination maps. To measure improvement, we calculated the Pearson correlation between estimated and actual log-rates across 100-kb windows and resampled the 120 paired regions 20,000 times. Recovery in bottlenecked populations improved as their effective size increased, and using paired large populations was more beneficial as the bottleneck became more severe (main-text Fig. 3C, D; SI Appendix, Fig. S5A).

For the empirical analysis in main-text Fig. 3A, B, we inferred maps across the first 40 Mb of chromosome 1 from 33 wolves and 67 domestic dogs, both scored against the Campbell dog pedigree map (11, 16). We estimated uncertainty by resampling eight contiguous 5-Mb blocks. The wolf-minus-dog differences were 0.051 (95% interval −0.039–0.179) at 100 kb and 0.055 (−0.125–0.268) at 1 Mb. Restricting the dog data to 42 Chinese village dogs gave differences of 0.043 (−0.049–0.160) and 0.038 (−0.152–0.186), respectively. For the LD-decay diagnostic, we randomly retained 33 dogs in each of 20 replicates, sampled 40,000 SNP pairs per distance bin in each dog replicate and 200,000 pairs per bin in wolves, and retained SNPs with minor-allele frequency 0.05–0.95. This dosage-$r^2$ diagnostic is separate from correlation between an inferred map and the pedigree map.

We expanded our inference to CanFam3 chromosomes 1–5 using 33 wolves and 42 village dogs. At 100 kb, raw-rate correlations for wolves were 0.534, 0.359, 0.365, 0.260, and 0.404, compared with 0.552, 0.330, 0.284, 0.162, and 0.316 for dogs. The chromosome means were 0.384 and 0.329, and the paired difference was 0.056 (chromosome-bootstrap 95% interval 0.012–0.091). The corresponding log-rate means were 0.318 and 0.271, a difference of 0.047 (0.028–0.065). The mean raw-rate differences increased from 0.068 at 200 kb to 0.096 at 500 kb and 0.118 at 1 Mb.

The strict marker-matching analysis used complete biallelic calls and 20 deterministic seeds. Each replicate randomly retained 33 of 42 village dogs, applied a minor-allele-frequency threshold of 0.05 separately in each population, and then retained the same number of variants in every 100-kb by 0.05-MAF. This yielded 248,304–250,635 variants in each matched population. After rerunning the frozen canid checkpoint, median 100-kb raw-rate correlation was 0.551 from wolves and 0.475 from dogs. The median paired gain was 0.077; a 10,000-draw interval that jointly selected a panel replicate and resampled 5-Mb blocks was 0.019–0.147. The corresponding log-Pearson gain was 0.028 (−0.022–0.093), and the Spearman gain was 0.041 (−0.016–0.104).A sensitivity analysis in which missing calls were imputed as the reference allele yielded 418,561–422,303 matching SNPs per population. Raw-rate gain was 0.026 (−0.013–0.166), log-Pearson gain was 0.051 (−0.001–0.126), and Spearman gain was 0.068 (0.005–0.143). Hence, the direction of the wolf–dog comparison was basically unaffected regardless of how missing calls were handled, yet the maginuted dependent on the used metric.

With respect to the analysis of chromosome 1, 1,227 100-kb windows were mapped to GC content in canFam3 along with 3,483 unique RefSeq TSSs. We performed a joint regression of standardized log rate on standardized GC, $\log(1 + \text{TSS count})$, and $\log(1 + \text{SNP count})$.

The adjusted GC coefficients were 0.165 (5-Mb block-bootstrap 95% interval 0.000–0.322) for the Campbell map, 0.185 (−0.073–0.398) for the wolf LD map, and 0.303 (0.130–0.477) for the dog LD map. The adjusted TSS-density coefficients were −0.101 (−0.219–0.027), −0.029 (−0.196–0.139), and 0.011 (−0.122–0.116), respectively. Although the unadjusted dog map had a promoter-density-quartile rate ratio of 1.269 (1.070–1.498), the adjusted TSS result shows that this enrichment was not separable from GC and SNP density at 100-kb resolution. We did not test a PRDM9 motif because PRDM9 is not functional in canids (35–37).

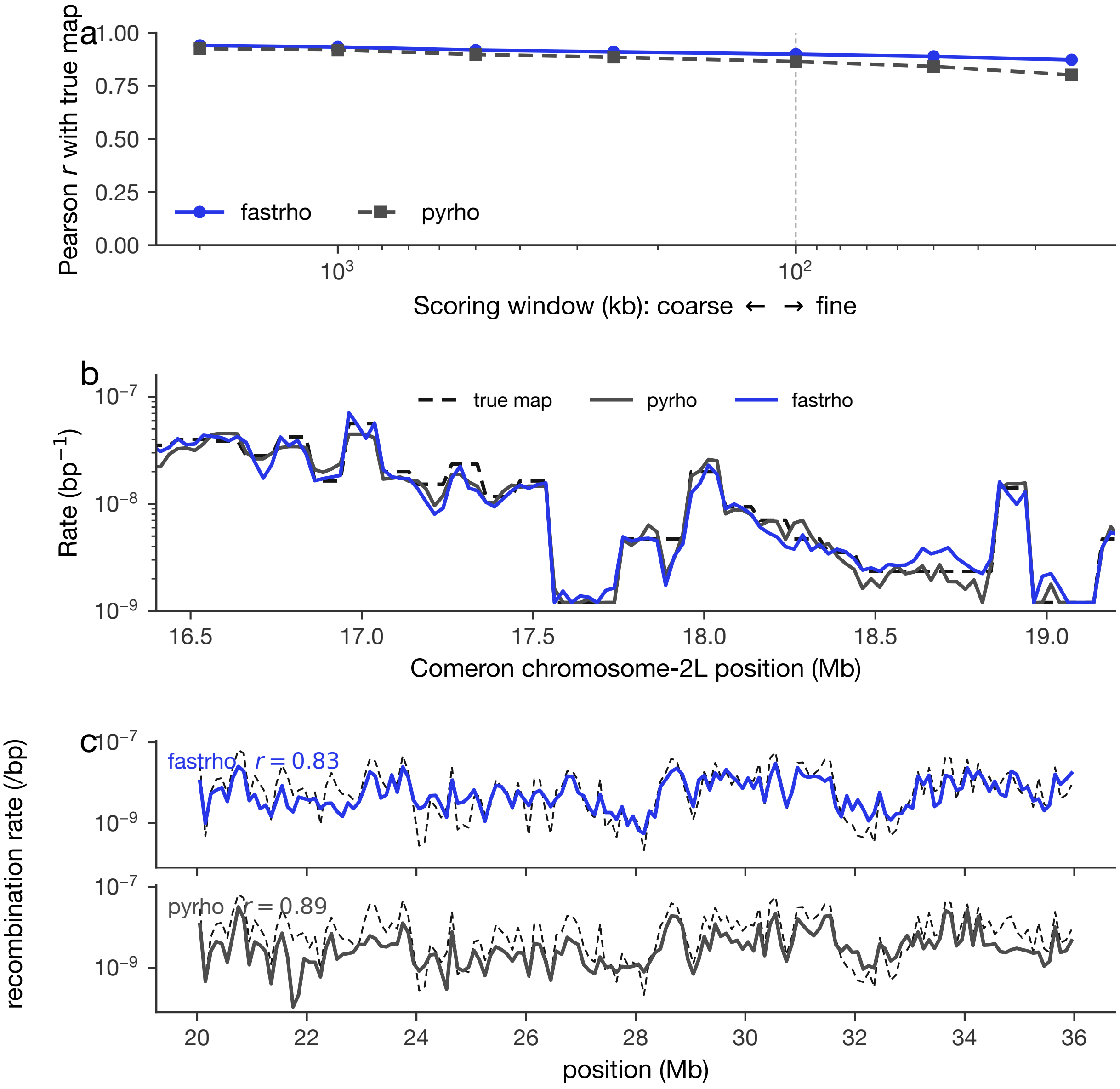


**Figure S3: Resolution and real-map detail.** (a) Pearson correlation for `fastrho` and `pyrho` with exact maps as the reporting grid changes from 2 Mb to 25 kb. (b) A Comeron chromosome-2L region showing their similar inferences. (c) CEU `fastrho` and `pyrho` profiles over the same dashed HapMap reference.

#### *Arabidopsis* selfing

For the selfing analysis, we used 156 south-Swedish *A. thaliana* haplotypes and aligned 100-kb windows across all five chromosomes. Following Nordborg (38), the selfing-aware model represented selfing rate $s$ using

$$F = \frac{s}{2-s}, \qquad N_{e,\text{eff}} = \frac{N_e}{1+F}, \qquad r_{\text{eff}} = r(1-F).$$

For `pyrho`, we kept the optimization settings fixed and rebuilt the lookup table under constant population size, the `stdpopsim` African2Epoch expansion, and the African3Epoch bottleneck; all three histories left chromosome-1 correlations near −0.35. We compared the inferred maps with the independent Rowan map, which contains 17,077 crossovers from 2,182 Col–Ler $F_2$ individuals (10) (main-text Fig. 3E–G). Missing calls, genotype error, polarization error, moderate population structure, and population expansion each changed exact-map recovery by no more than approximately 0.12, and none reproduced the empirical difference between random-mating and selfing-aware inference (SI Appendix, Fig. S5B).

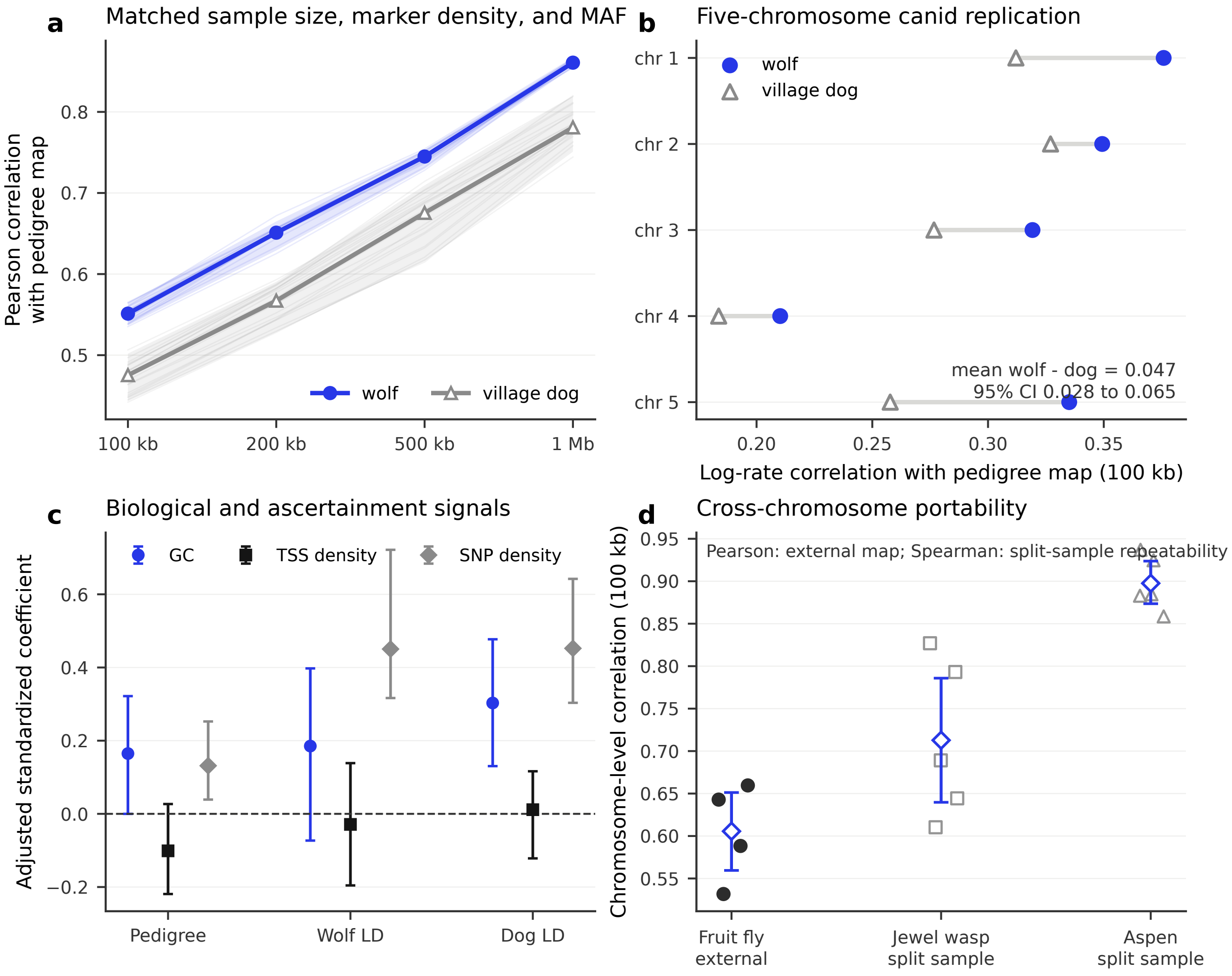


**Figure S4: Analyses addressing demographic and marker-ascertainment concerns.** (a) Pearson correlation with the Campbell pedigree map after choosing 33 individuals per population and the complete-call variant count within each 100-kb by 0.05-MAF bin. Lines represent 20 samples, thick lines are medians, and shading is the 2.5th–97.5th percentile range. (b) Log-rate correlation with the pedigree map on CanFam3 chromosomes 1–5; gray segments pair populations within chromosome and the label is a chromosome-bootstrap confidence interval for the mean difference. (c) Standardized regression coefficients for a log-rate model with GC content, TSS density, and SNP density in 100-kb windows of chromosome 1; intervals are 5-Mb block-bootstrap 95% intervals. (d) Chromosome-level replication of the portability screen. Fruit fly points are Pearson correlations with an independent Comeron map; jewel wasp and aspen points are split-sample Spearman correlations and hence test repeatability. Diamonds are chromosome averages and intervals are chromosome-bootstrap 95% intervals.

#### Cross-species comparison

To further evaluate our method, we used ten publicly available datasets spanning mammals, insects, and plants. The accessions, sample sizes, analyzed chromosomes, and sources are listed in Table S2 (5–8, 20–26). We analyzed one population and one representative chromosome per species, except for *Arabidopsis*, for which results were averaged across five chromosomes. All species used the same fixed model except *Arabidopsis*, which used the selfing model. Retained panels had at least 40 haplotypes, 8,000 segregating SNPs, 40 scored 100-kb windows, and a variable inferred recombination map. Species with an external map required Pearson $r \geq 0.25$; those without one required split-sample correlation $\rho \geq 0.50$ and agreement with pyrho of at least $r = 0.50$ after rounding.

Seven panels have straightforward population-sample interpretations. Donkey and Chinese chestnut combine sampling localities, whereas the jewel-wasp panel consists of 34 lines derived from a single outbred population and subsequently inbred (22). These three panels are therefore marked.

External-map comparisons used Pearson correlations to compare recombination rates across aligned 100-kb windows. In the

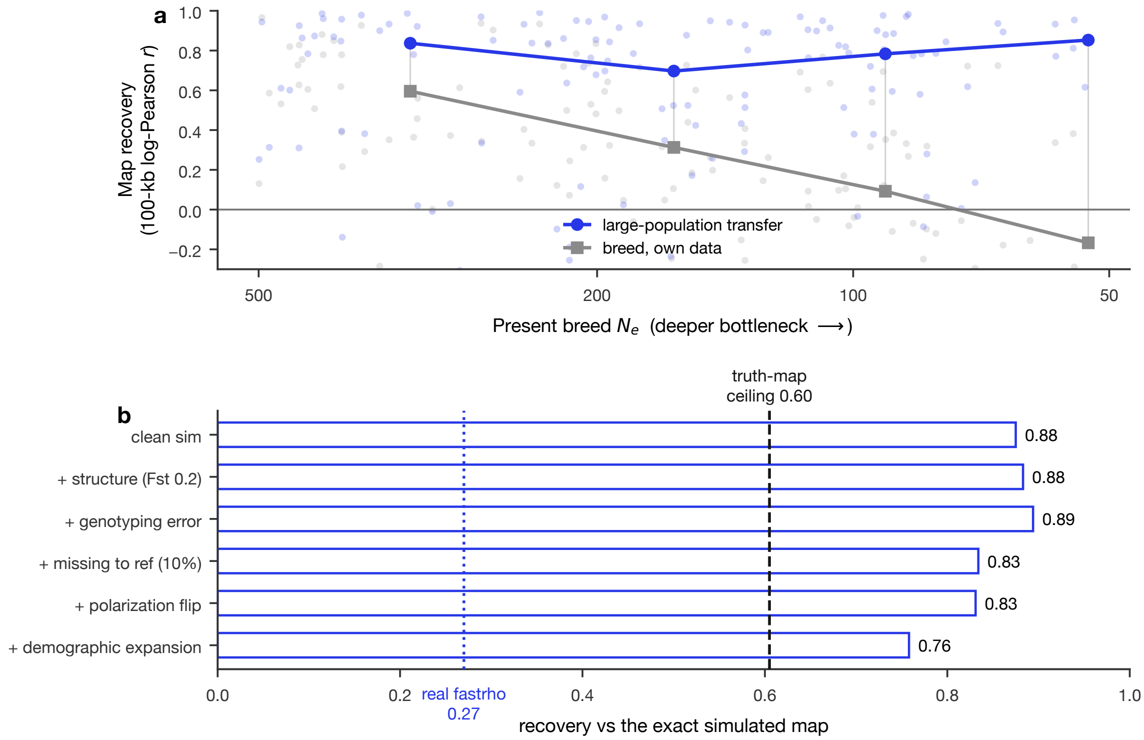


**Figure S5: Demographic and mating-system limits on recovery.** (a) Canid map rescue increases with bottleneck severity. The points represent the 120 pairs of simulated populations ranked by 100 kb log-Pearson correlation, while the lines are the medians for the present $N_e$ bins, and the vertical connecting lines denote the differences between these medians. Gray lines are the estimates obtained based on the LD in the bottlenecked population itself, while the cobalt lines are based on the large population used to construct the pairs. (b) One confounder cannot explain the selfing difference. We demonstrate the correlations due to each confounding factor, such as structure or genotyping error, via simulations. The shared-map canid trajectories, recovery ladder, and chromosome-level empirical results appear in the main text Fig. 3 and are not repeated.

case where there were no external maps available, $\rho$ represents the Spearman correlation between the recombination maps derived from separate halves of the sample and so reflects repeatability and not accuracy. The Pearson correlation with `pyrho`, $r_{\texttt{pyrho}}$, similarly entails comparing two different LD estimates based on the same data.

**Table S2:** Sources and map comparisons for the ten species. Here, $n$ is the number of diploid individuals, except for DGRP and jewel wasp, where it is the number of inbred lines; Chr. is the analyzed chromosome or arm. Three species report Pearson $r$ against an external map. The other seven report split-sample Spearman reproducibility $\rho$ and Pearson agreement with `pyrho`, $r_{\texttt{pyrho}}$.

| Common name / *species* | Data source (accession) | $n$ | Chr. | Recovery |
|---|---|---|---|---|
| **Mammals** | | | | |
| Human / *Homo sapiens* | IGSR 1000G `20130502`, CEU | 99 | 2 | $r$=+0.87 (HapMap) |
| Cattle / *B. taurus* × *B. indicus* | EBI NextGen `UGBT` (Uganda) | 25 | 1 | $\rho$=0.59; $r_{\texttt{pyrho}}$=0.60 |
| Sheep / *Ovis aries* | EBI NextGen `IROA` (Iran) | 20 | 1 | $\rho$=0.73; $r_{\texttt{pyrho}}$=0.75 |
| Goat / *Capra hircus* | EBI NextGen `IRCH` (Iran) | 20 | 1 | $\rho$=0.72; $r_{\texttt{pyrho}}$=0.82 |
| Donkey† / *Equus asinus* | Todd 2022 (Chinese cohort) | 51 | CM027690.1 | $\rho$=0.86; $r_{\texttt{pyrho}}$=0.73 |
| **Insects** | | | | |
| Fruit fly / *D. melanogaster* | DGRP inbred lines | 205 lines | 2L | $r$=+0.49 (Comeron) |
| Jewel wasp† / *N. vitripennis* | EVA `PRJEB33514` | 34 lines | NC_015868.2 | $\rho$=0.72; $r_{\texttt{pyrho}}$=0.50 |
| **Plants** | | | | |
| Thale cress / *A. thaliana* | 1001 Genomes | 78 | (1–5) | $r$=+0.27 (Salomé) |
| Aspen / *Populus tremula* | SwAsp (`PRJEB79788`) | 99 | chr1 | $\rho$=0.94; $r_{\texttt{pyrho}}$=0.64 |
| Chinese chestnut† / *C. mollissima* | EVA `PRJEB87510` | 97 | Chr1 | $\rho$=0.92; $r_{\texttt{pyrho}}$=0.57 |

For the seven species without external fine-scale maps, split-sample correlations ranged from 0.59 to 0.94. Agreement with `pyrho` was between 0.50 and 0.82 (Fig. S6).

For the chromosome-level follow-up analysis, we kept all compatible chromosomes that passed the same input criteria for three sets of data: arms 2L, 2R, 3L, and 3R of *D. melanogaster*, and five chromosomes each for *N. vitripennis* and *P. tremula*. For each chromosome, the fixed checkpoint was run on the full dataset and on two different half-sample sets. The correlation analysis was done at 100, 200, 500, and 1,000 kb. Chromosomes were resampled 10,000 times to get an uncertainty estimate. At 100 kb, the correlation values for the four *Drosophila* arms with respect to the Comeron map were 0.588, 0.643, 0.660, and 0.532, with a chromosome average of 0.606 (95% interval 0.559–0.651). For the jewel-wasp chromosomes, the split-sample Spearman correlation values ranged from 0.610 to 0.827, while those for the five aspen chromosomes ranged from 0.859 to 0.937, with averages of 0.713 (0.640–0.786) and 0.898 (0.874–0.924), respectively (Fig. S4D).

#### Ag3.0 populations and variant processing

The empirical atlas was based on the Ag3.0 phased-haplotype release. The 13 predetermined populations consisted of six *Anopheles gambiae*, four *Anopheles coluzzii*, and three *Anopheles arabiensis* populations. We sampled exactly 40 diploid individuals without replacement from each population using seed 2026, giving 520 unique mosquitoes and 80 haplotypes per population; the same individuals were used on all five chromosome arms. Sites were single-contig, biallelic, segregating, and complete, and VCF positions were converted from one-based coordinates to zero-based, half-open BED intervals. (1)

Identifiers for the populations, numbers of samples, numbers of SNPs, and frequencies of the 2La arrangements are given in Table S3.

**Table S3:** Ag3.0 populations included in the atlas. Map panels contained 80 haplotypes from 40 diploid mosquitoes. $H_{40}$ is expected 2La heterokaryotype frequency in that map panel; $n_{\text{full}}$ and $H_{\text{full}}$ use all eligible Ag3 tag-SNP calls and were used in the primary 2La analysis.

| Population | Species | Country | Haplotypes | SNPs | $H_{40}$ | $n_{\text{full}}$ | $H_{\text{full}}$ |
|---|---|---|---|---|---|---|---|
| arab_MW | *A. arabiensis* | Malawi | 80 | 7,014,160 | 0.004 | 41 | 0.004 |
| arab_TZ | *A. arabiensis* | Tanzania | 80 | 7,873,436 | 0.003 | 225 | 0.003 |
| arab_UG | *A. arabiensis* | Uganda | 80 | 7,939,204 | 0.004 | 82 | 0.004 |
| colu_BF | *A. coluzzii* | Burkina Faso | 80 | 8,580,694 | 0.026 | 135 | 0.031 |
| colu_CI | *A. coluzzii* | Cote d'Ivoire | 80 | 6,930,629 | 0.480 | 80 | 0.462 |
| colu_GM | *A. coluzzii* | Gambia | 80 | 8,879,083 | 0.410 | 200 | 0.449 |
| colu_ML | *A. coluzzii* | Mali | 80 | 9,065,121 | 0.030 | 90 | 0.036 |
| gamb_BF | *A. gambiae* | Burkina Faso | 80 | 9,700,565 | 0.183 | 158 | 0.169 |
| gamb_CM | *A. gambiae* | Cameroon | 80 | 9,775,041 | 0.439 | 416 | 0.478 |

| Population | Species | Country | Haplotypes | SNPs | $H_{40}$ | $n_{\text{full}}$ | $H_{\text{full}}$ |
|---|---|---|---|---|---|---|---|
| gamb_GA | *A. gambiae* | Gabon | 80 | 4,953,910 | 0.049 | 69 | 0.070 |
| gamb_GN | *A. gambiae* | Guinea | 80 | 9,526,350 | 0.480 | 125 | 0.482 |
| gamb_ML | *A. gambiae* | Mali | 80 | 9,441,137 | 0.201 | 131 | 0.216 |
| gamb_UG | *A. gambiae* | Uganda | 80 | 8,587,341 | 0.500 | 207 | 0.492 |

#### Ag3 recombination atlas

The adjacent-SNP predictions were categorized based on physical distance in non-overlapping 50-kb windows on arms 2R, 2L, 3R, 3L, and X. For each species track, the median values are shown in bold, while the different populations are represented using thin lines. Differences in vertical scale between the different species are not considered to represent differences in crossover rates, since $r$ is conditional on the population-specific value of $N_e$.

#### Ag3 pedigree crossover calling and map comparison

The Ag3.0 pedigree data contain 297 samples from 15 laboratory crosses. Because the public metadata list their taxon as unassigned, we refer to them collectively as *An. gambiae*-complex crosses without assigning species to individual parents. The primary direct map used five crosses held out during construction of the Ag3 site filter and was compared with the inferred map summarized across the ten *Anopheles gambiae*/*Anopheles coluzzii* atlas populations; all 15 crosses provided a sensitivity analysis.

Crossovers on arms 2R, 2L, 3R, and 3L were identified from changes in transmitted parental haplotypes across quality-filtered markers. A switch required at least three called markers on each side and a breakpoint interval no longer than 1 Mb. The caller was tested with simulations matched to the observed marker structure and by inserting synthetic crossovers into observed transmission backgrounds. The sensitivity analyses involved changes in quality of markers, event classification, breakpoint resolution, family inclusion and genomic region. The initial analysis was carried out with normalization for each chromosome arm, with correlation performed using 5 Mb window sizes. The second level analysis involved 2 Mb windows.

#### Local `pyrho` comparison

We did select ten diploids from one population per species for a comparison on 3R:6–14 Mb and gave both estimators the same 20 haplotypes. `pyrho` used a constant-size lookup table based on Watterson's estimate, mutation rate $3.5 \times 10^{-9}$, optimization window 50, ploidy one, and block penalty 25.

#### Positive control analysis for Ag3 2La

The 2La region is 20.524–42.166 Mb long in AgamP4. The ratio of median positive rates within the inversion to median positive rates on 2L outside the inversion was calculated separately for each population. Suppression depth was defined as one minus this ratio. The expected frequency of heterokaryotypes was estimated based on all the Ag3 arrangements called for each population, which ranged from 41 to 416 mosquitoes per population (total of 1,959).

#### Resistance gene regions and their matches

The list of 15 literature-reported AgamP4 regions with phenotype-associated markers or uniquely mappable *Anopheles gambiae* resistance genes was used. Adjacent genes mapped in the same window were combined into a single region. Conflicting gene families, microbes, inversions, and regions only available in *Anopheles funestus* were not considered. Details about the included regions are listed in Table S4. (39, 40)

**Table S4:** Definitions of the 15 insecticide-resistance regions. Positions are AgamP4 coordinates.

| Region | Arm | Position (Mb) | Mechanism | Inclusion basis |
|---|---|---|---|---|
| *Vgsc/kdr* | 2L | 2.390 | Target site | Phenotype-associated resistance locus |
| *Rdl* | 2L | 25.400 | Target site | Phenotype-associated resistance locus |
| *Ace1* | 2R | 3.500 | Target site | Phenotype-associated resistance locus |
| *Gste2* | 3R | 28.600 | Metabolic detoxification | Phenotype-associated resistance locus |
| *Cyp6aa/Cyp6p* | 2R | 28.500 | Metabolic detoxification | Phenotype-associated resistance cluster |
| *Cyp9k1* | X | 15.240 | Metabolic detoxification | Phenotype-associated resistance locus |

| Region | Arm | Position (Mb) | Mechanism | Inclusion basis |
|---|---|---|---|---|
| *Cyp4j5* | 2L | 25.635 | Metabolic detoxification | Phenotype-associated resistance locus |
| *Coeae1d* | 2L | 20.288 | Metabolic detoxification | Phenotype-associated resistance locus |
| *Cyp6m2* | 3R | 6.930 | Metabolic detoxification | Named mechanism locus |
| *SAP2* | 3R | 4.869 | Insecticide sequestration | Named mechanism locus |
| *Cyp4g16* | X | 22.942 | Cuticular resistance | Named mechanism locus |
| *Cyp4g17* | X | 16.620 | Cuticular resistance | Named mechanism locus |
| *Gstd1* | 2R | 50.999 | Metabolic detoxification | Named mechanism locus |
| *Maf-S* | 3L | 2.786 | Metabolic regulation | Named mechanism locus |
| *D7r2/D7r4* | 3R | 8.559 | Insecticide sequestration | Adjacent named genes represented as one region |

The focal rate for each region was defined as the median rate in a 0.15-Mb interval. The controls included were on the same arm, at least 0.50 Mb away from all focal regions, and matched on empirical percentiles of relative chromosome position, log SNP density, nucleotide diversity, and H12. The union of all 15 focal regions was removed from the pool of controls for all loci, and up to 75 controls were selected for each locus. H12 was computed using a maximum of 128 deterministically sampled SNPs in each 0.30-Mb interval. We drew 5,000 permutation samples for each population and took the geometric mean across all loci as our population statistic.

#### Processing and scoring of the *Arabis* hybrid cross map

The *Arabis* analysis used ENA projects PRJEB39992, PRJEB33482, and PRJEB89863 and the linkage map of Rahnamae et al. (18), and species identification followed Dittberner et al.; see Supplementary Table S2 (19). We aligned the reads and linkage markers against the chromosome-level assembly of *A. nemorensis* GCA_976985625.1. The linkage map was constructed from 742 $F_2$ hybrids resulting from reciprocal interspecific crosses. We collapsed duplicate positions to their median centimorgan coordinates, oriented chromosomes by physical–genetic rank correlation, and made genetic positions nondecreasing by isotonic regression.

The populations included 12 *A. nemorensis* and 25 *A. sagittata* genomes. Within each species, we removed sites with missing or heterozygous calls and kept one allele per predominantly selfing accession. We inferred separate maps for the two species and compared both with the same $F_2$ map. We first applied a model trained with 50–200 selfing lines without any *Arabis*-specific training. We divided each map by its chromosome mean and combined the two inferred maps using their geometric mean. We used Spearman correlation across pooled 2-Mb windows as the primary statistic. As sensitivity analyzes, we used 1- and 5-Mb windows, excluded distorted chromosomes 4 and 7 or the two $F_2$ parents, omitted each chromosome in turn, resampled chromosomes to estimate confidence intervals, and circularly shifted windows within chromosomes to construct a null distribution.

We developed two follow-up approaches after the baseline comparison. For the sample-size analysis, we trained five models on 8,000 regions containing 8–50 lines. For the structured-selfing experiment, we simulated 16,000 training, 1,600 validation, and 1,600 test regions assuming panmictic mating, weak island structure. Selfing rate, migration, split time, population size, mutation rate, recombination rate, map shape, and panel size were varied. We selected seven models via a simulated validation procedure and evaluated each on independent simulated maps before applying it to empirical data. The $F_2$ map was used for scoring results only.

Our quality control procedures allowed us to retain all 102 windows after checking SNP count, allele frequency, LD, SNP gaps, border processing, and prediction variance.

We found no association between the sharp peaks in the chromosome-1 and chromosome-3 windows and either sparse data or shared processing boundaries, and all seven structured-selfing models supported their elevated rates. Figure S8D summarizes cumulative maps across chromosomes, and Fig. S8E groups windows by the corresponding $F_2$ rate for display.

At baseline, correlations had the same direction at 1, 2, and 5 Mb, and the 2-Mb consensus remained positive when each chromosome was omitted ($r_s$ = 0.157–0.408). Excluding distorted chromosomes 4 and 7 increased the correlations to 0.443 for *A. sagittata*, 0.425 for the consensus, and 0.236 for *A. nemorensis*. Matching the simulated panel size increased the *A. sagittata* and consensus correlations from 0.330 to 0.519 and from 0.305 to 0.378, respectively, but changed the *A. nemorensis* correlation from 0.158 to −0.021. Requiring the minor allele in at least two *A. nemorensis* accessions left its correlation near zero ($r_s$ = 0.028). Equal-sized *A. sagittata* subsets remained correlated with the cross map (median $r_s$ = 0.592), whereas leave-one-out and geographically restricted *A. nemorensis* panels remained discordant (medians −0.037 and −0.206). Panel size and rare variants therefore did not explain the species difference.

The inference produced 2-Mb correlations of $r_s$ = 0.270 for *A. nemorensis* (chromosome-bootstrap 95% interval −0.034–0.545; spatial-shift $P$ = 0.0464), 0.657 for *A. sagittata* (interval 0.496–0.803; $P$ = 0.00010), and 0.536 for the consensus (interval 0.306–0.728; $P$ = 0.00070). Removing the $F_2$ parental accessions increased the *A. nemorensis* and consensus correlations to 0.354

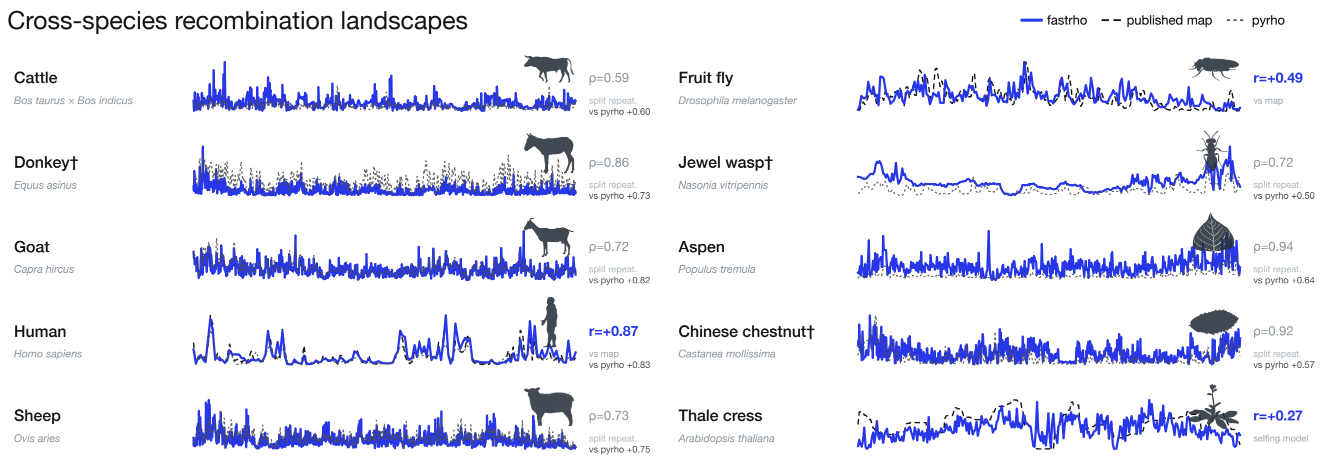


**Figure S6: Cross-species recombination-map comparison.** One representative chromosome is shown for each species. The cobalt curves represent `fastrho` estimates for unphased genotypes, whereas dashed curves represent published maps whenever available. The dotted gray curves indicate `pyrho` estimates for the same set of genotypes. The maps are normalized within species, thus emphasizing their shapes. Pearson $r$ values compare `fastrho` with the published map at 100 kb, while the gray values of $\rho$ represent comparisons between independent samples. Panels with limitations are marked. Silhouettes are from PhyloPic.

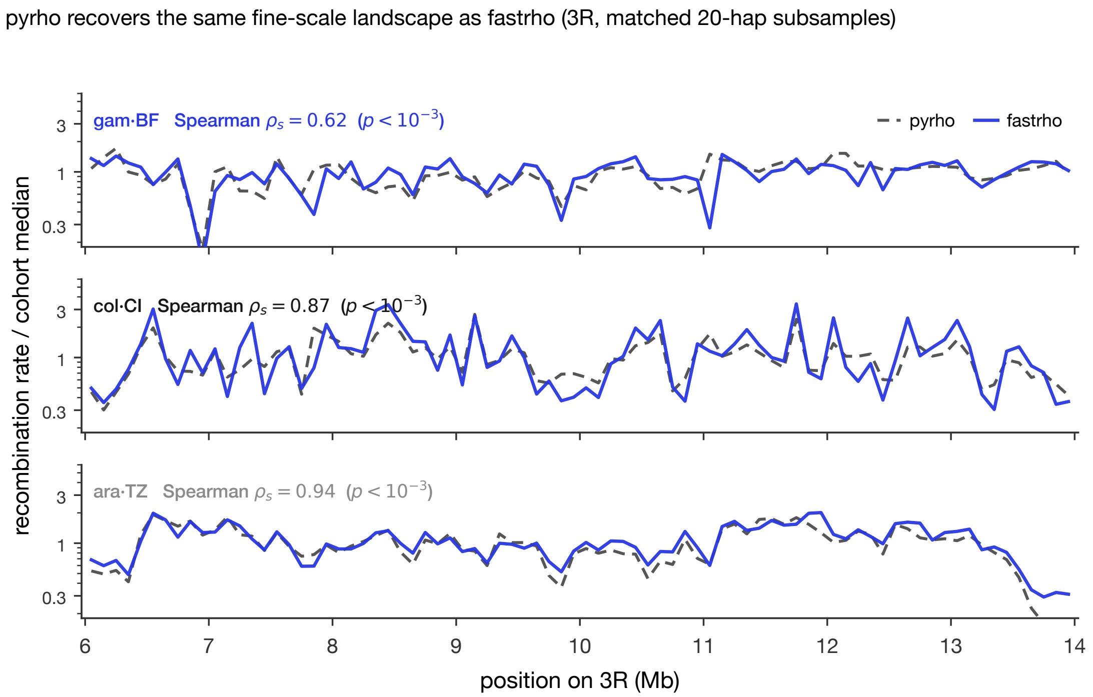


**Figure S7: Local `pyrho` comparison of selected Ag3 map segments.** Solid cobalt `fastrho` and dashed gray `pyrho` profiles use matched 20-haplotype subsamples from one population per species along chromosome arm 3R (6–14 Mb), summarized in 100-kb windows and divided by each population median. Annotations give Spearman correlations across windows.

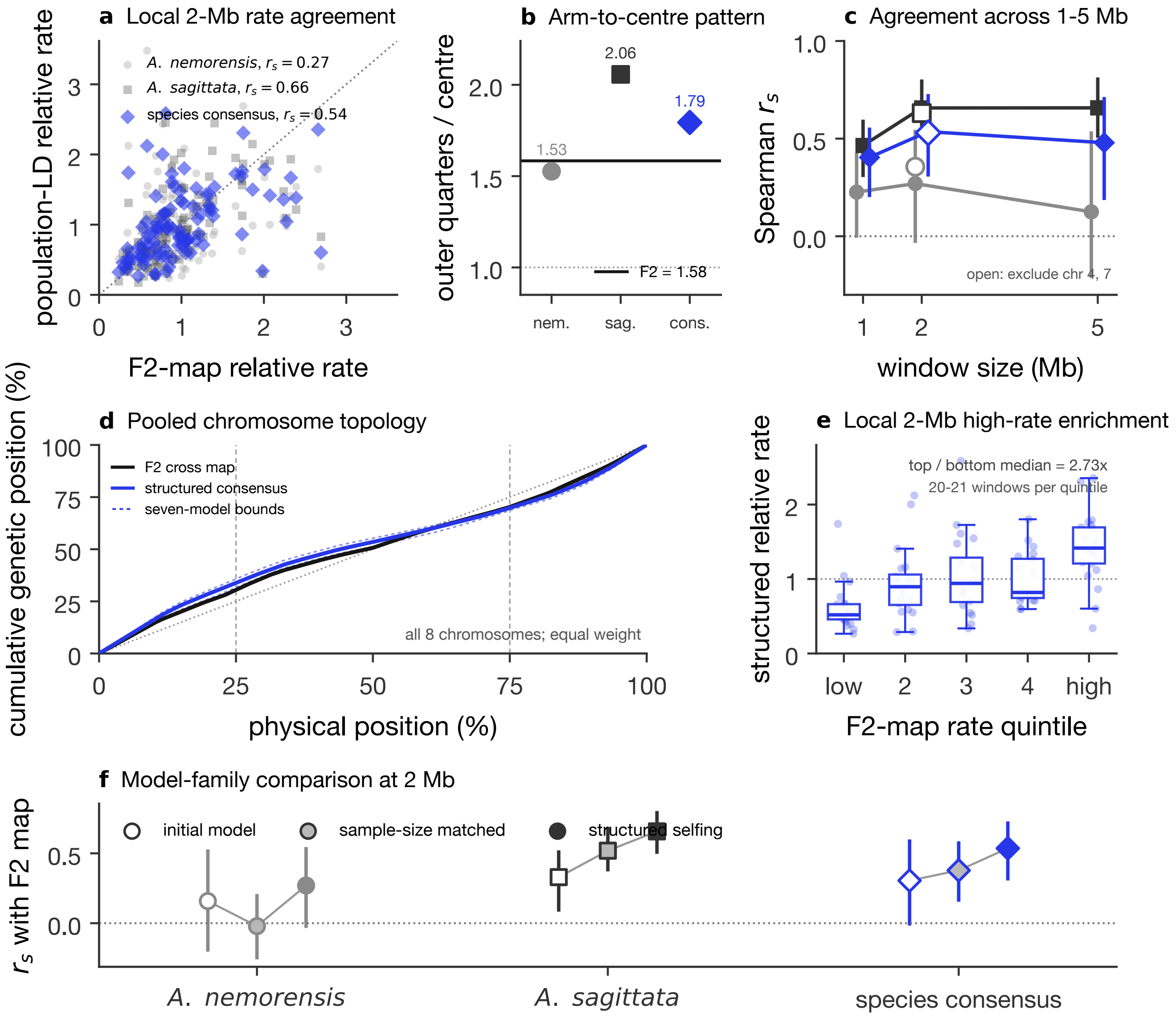


**Figure S8: Comparison with the *Arabis* $F_2$ linkage map.** The 742 offspring form one interspecific map. Panels (a)–(e) show results from the structured-selfing ensemble. We inferred recombination maps separately from 12 *A. nemorensis* and 25 *A. sagittata* genomes on the *A. nemorensis* assembly. (a) Rates in 102 2-Mb windows. The combined estimate is the geometric mean of the chromosome-normalized species maps. (b) Rate in the outer chromosome quarters relative to the central half. (c) Rank correlation at 1, 2, and 5 Mb; open symbols exclude distorted chromosomes 4 and 7. Lines are 95% chromosome-bootstrap intervals. (d) Mean cumulative maps across eight chromosomes. (e) Inferred rates grouped by the corresponding fifth of the $F_2$-rate distribution. (f) Initial, sample-size-matched, and structured-selfing correlations at 2 Mb. Follow-up models were selected using simulations but designed after the initial empirical comparison.

and 0.578, respectively, while leaving the *A. sagittata* value unchanged. Across all 102 windows, the median consensus rate was 2.73-fold higher in the highest than in the lowest $F_2$-rate fifth. The structured consensus and $F_2$ cumulative maps differed by 3.4, 2.7, and 0.5 percentage points at 25, 50, and 75% of chromosome length (Fig. S8D, E). However, these simulations were designed after the baseline comparison, did not identify a distinct demographic history, and showed scale sensitivity for *A. nemorensis* ($r_s$ = 0.226 at 1 Mb and 0.125 at 5 Mb).

#### Trained models used in the study

We trained both general and regime-specific `fastrho` models on population-genetics regions simulated using different types of inputs and biological settings, being explicit that empirical maps were not our optimization objectives. Table S5 lists the training data and figures associated with each checkpoint and/or ensemble, while Table S9 shows the simulation design and parameter values used to generate those regions. Boldface names refer to models that are publicly available for further compatible studies, and other models are shown for reproduction purposes.

**Table S5:** Trained `fastrho` models used in the manuscript and Supporting Information.

| Name | Description and training data | Figures used |
|---|---|---|
| **`domain-randomized-v1`** | Multi-view model trained on 15,000 simulated regions from the broad demographic prior, with each region represented as phased haplotypes, unphased dosages, and folded unpolarized dosages. | Main Fig. 1; SI Figs. S2 and S6 |
| **`base-v1`** | Single-view model trained on phased, polarized haplotypes from the same broad demographic prior of constant, sawtooth, island, and bottleneck histories. | Main Fig. 2; SI Figs. S1, S2, and S3 |
| **`composite-ld-v1`** | Single-view specialist trained by re-featurizing the 15,000 base-prior simulations as folded, unphased diploid genotypes. | Main Fig. 6; SI Fig. S2 |
| **`high-ne-v1`** | Phased specialist trained on 15,000 simulated high-diversity, high-$N_e$ regions representing mosquito and other dipteran regimes. | Main Figs. 5 and 4; SI Fig. S7 |
| **`selfing-v1`** | Phased specialist trained on 4,000 simulated regions with 50–200 predominantly selfing lines and selfing-adjusted ancestry and recombination. | Main Fig. 3E–G; SI Figs. S5B, S6, and S8F |
| **`dog-bottleneck-v1`** | Folded composite-LD specialist trained on 15,000 simulated unphased canine panels with village-dog and severe breed-like bottleneck histories. | Main Fig. 3A, C, D; SI Fig. S5A |
| `arabis-smalln-ensemble` | Five models, each trained on 8,000 simulated selfing regions containing 8–50 lines, for the sample-size follow-up. | SI Fig. S8F |
| `arabis-structured-ensemble` | Seven models trained on 16,000 simulated regions spanning panmixia, weak island structure, and the two empirical sampling layouts. | SI Fig. S8A–F |
| `canid-structure-paper-analysis` | One checkpoint fine-tuned on 1,200 simulated regions with 8–20 diploids under panmixia and two- or three-deme island models. | SI full-chromosome sensitivity analysis |

### SI Tables

**Table S6:** Benchmark Pearson correlations at 100- and 25-kb reporting scales for `fastrho` and `pyrho`. In the scenario column, $n$ is the number of sampled haplotypes. The bottleneck and expansion rows use the paired `fastrho` reference and the `pyrho` generating-history lookup table; see Table S8 for both `pyrho` specifications. All paired values use the same VCFs, true maps, and valid windows.

| config | Pearson @100kb | | Pearson @25kb | |
|---|---|---|---|---|
| | `fastrho` | `pyrho` | `fastrho` | `pyrho` |
| const_n20 | 0.893 | 0.875 | 0.882 | 0.840 |
| const_n40 | 0.911 | 0.914 | 0.892 | 0.864 |
| real_hapmap | 0.873 | 0.846 | 0.833 | 0.793 |
| real_decode | 0.921 | 0.877 | 0.886 | 0.807 |
| bottleneck_n20 | 0.833 | 0.732 | 0.776 | 0.619 |
| expansion_n20 | 0.682 | 0.618 | 0.604 | 0.444 |
| real_dog | 0.892 | 0.868 | 0.834 | 0.788 |
| const_n100 | 0.952 | 0.970 | 0.941 | 0.952 |

**Table S7:** Window-matched Pearson correlations at the ReLERNN window scale for main-text Fig. 2A (lower subpanel). Each scenario contains 20 prespecified 10-Mb regions with one constant rate per region. The exact truth, fastrho, and pyrho predictions were averaged within each complete ReLERNN output interval. "Matched" denotes the generating demographic history; "constant" is a deliberate misspecification applied to the identical bottleneck or expansion inputs. "Windows" is the number of complete intervals shared by all methods.

| Scenario | History | Median window (kb) | Windows | fastrho | pyrho | ReLERNN |
|---|---|---|---|---|---|---|
| Constant | matched | 740 | 259 | 0.964 | 0.956 | 0.946 |
| Bottleneck | matched | 1,649 | 112 | 0.959 | 0.943 | 0.898 |
| | constant | 1,649 | 112 | 0.959 | 0.892 | 0.854 |
| Expansion | matched | 2,853 | 60 | 0.942 | 0.961 | 0.847 |
| | constant | 2,853 | 60 | 0.942 | 0.967 | 0.772 |
| deCODE | matched | 734 | 273 | 0.967 | 0.966 | 0.931 |
| HapMap | matched | 743 | 262 | 0.978 | 0.969 | 0.937 |
| Dog | matched | 2,282 | 80 | 0.966 | 0.990 | 0.926 |

**Table S8:** Effect of demographic specification in the paired 25-kb benchmark. The fixed fastrho estimator was trained once over a broad demographic prior and was not tuned for either history. Constant and matched refer to the history used to construct the pyrho lookup table. Estimated/true median indicates absolute-rate calibration, where one is perfect.

| Scenario | Method | Demographic specification | Pearson $r$ | Spearman $\rho$ | Median estimated/true |
|---|---|---|---|---|---|
| Bottleneck | fastrho | fixed broad prior | 0.776 | 0.512 | 0.891 |
| | pyrho | constant | 0.563 | 0.453 | 0.283 |
| | pyrho | matched | 0.619 | 0.467 | 0.788 |
| Expansion | fastrho | fixed broad prior | 0.604 | 0.459 | 0.687 |
| | pyrho | constant | 0.447 | 0.426 | 0.364 |
| | pyrho | matched | 0.444 | 0.436 | 0.829 |

**Table S9:** Simulation data and prior ranges for each released model and paper-specific follow-up. These ranges define the evaluated amortization priors; they do not guarantee accuracy for every dataset.

| Model or ensemble | Simulated training data | Prior and design |
|---|---|---|
| base-v1 | 15,000 1-Mb regions represented as phased, polarized haplotypes; 20–200 haplotypes per region. | $\mu = 3.2\times10^{-9}$–$2.0\times10^{-8}$; $\bar{r} = 2.0\times10^{-9}$–$2.0\times10^{-8}$; $N_e = 3.2\times10^{3}$–$6.3\times10^{4}$; constant, sawtooth, three-deme island, and bottleneck histories; GP, hotspot, or constant recombination maps. |
| domain-randomized-v1 | The same 15,000 broad-prior regions, each represented as phased haplotypes, unphased dosages, and folded unpolarized dosages. | The same demographic and map prior as base-v1; training randomized the input representation rather than the demographic history. |
| composite-ld-v1 | The same 15,000 broad-prior regions re-featurized as folded, unphased diploid genotypes. | The same demographic and map prior as base-v1; one composite-LD input view only. |
| high-ne-v1 | 15,000 phased, polarized high-diversity regions initialized at 30 kb and shortened when required for tractable simulation. | The broad prior extended to $N_e = 2\times10^{6}$ and mean recombination rates of $5.0\times10^{-8}$ for mosquito and other dipteran regimes. |
| selfing-v1 | 4,000 1-Mb phased regions containing 50–200 predominantly selfing lines, with one haplotype retained per line. | $\mu = 5\times10^{-9}$–$1.0\times10^{-8}$; census $N_e = 10^{5}$–$4\times10^{5}$; selfing rate $s = 0.90$–$0.99$; ancestry and recombination rescaled by the inbreeding coefficient. |
| dog-bottleneck-v1 | 15,000 folded, unphased canine regions containing 60–134 haplotypes. | $\mu = 2\times10^{-9}$–$6\times10^{-9}$; village-dog and breed-like histories with severe recent crashes, reaching $N_e = 50$. |
| arabis-smalln-ensemble | Five models, each trained on 8,000 1-Mb selfing regions with 8, 12, 16, 24, 25, 32, or 50 sampled lines. | $\mu = 10^{-8.3}$–$10^{-8.0}$; census $N_e = 10^{5}$–$10^{5.6}$; meiotic $\bar{r} = 10^{-8.0}$–$10^{-7.4}$; $s = 0.90$–$0.99$; GP or hotspot maps. |
| arabis-structured-ensemble | Seven models trained on 16,000 1-Mb regions, with 1,600 validation and 1,600 independent test regions; panel sizes were 8, 12, 16, 24, 25, 32, or 50. | 30% panmixia, 30% diffuse island structure, and 20% for each empirical sampling layout; census $N_e = 10^{4.7}$–$10^{5.7}$; $s = 0.70$–$0.997$; GP, hotspot, or pericentromeric maps. |
| canid-structure-paper-analysis | One checkpoint fine-tuned on 1,200 simulated regions with 8–20 diploid samples and evaluated on 240 independent regions. | $N_e = 15{,}000$–$80{,}000$; $\mu = 2\times10^{-9}$–$6\times10^{-9}$; 45% panmixia and otherwise two- or three-deme island models with migration $10^{-5.2}$–$10^{-3}$. |

[27] Graham Gower, Nathaniel S. Pope, Murillo F. Rodrigues, Silas Tittes, Linh N. Tran, Ornob Alam, Maria Izabel A. Cavassim, Peter D. Fields, Benjamin C. Haller, Xin Huang, Ben Jeffrey, Kevin Korfmann, Christopher C. Kyriazis, Jiseon Min, Inés Rebollo, Clara T. Rehmann, Scott T. Small, Chris C. R. Smith, Georgia Tsambos, Yan Wong, Yu Zhang, Christian D. Huber, Gregor Gorjanc, Aaron P. Ragsdale, Ilan Gronau, Ryan N. Gutenkunst, Jerome Kelleher, Kirk E. Lohmueller, Daniel R. Schrider, Peter L. Ralph, and Andrew D. Kern. Accessible, realistic genome simulation with selection using stdpopsim. *Molecular Biology and Evolution*, 42(11): msaf236, 2025. doi: 10.1093/molbev/msaf236.
